# Decoding the comprehensive substrate-specificity and evidence of altered site-specific collagen prolyl-3-hydroxylation, lysyl-hydroxylation, and lysyl O-glycosylation in P4ha1 and P4ha2 deleted mutant mice

**DOI:** 10.1101/2023.06.28.546985

**Authors:** Vivek Sarohi, Trayambak Basak

## Abstract

Collagens, the most abundant proteins in mammals, play pivotal roles in the maintenance of tissue structure, functions, cell-to-cell communication, cellular migration, behavior, and growth. Collagens are highly complex in structure due to the dynamic post-translational modifications (PTMs) such as hydroxylations (on prolines and lysine residues) and O-glycosylation (on hydroxylysines) enzymatically catalyzed during biosynthesis. The most prevalent modification in fibrillar collagens is prolyl 4-hydroxylation catalyzed by collagen prolyl 4-hydroxylases (C-P4hs). Prolyl 4-hydroxylation on collagens plays a critical role in collagen biosynthesis, thermostability, and cell-collagen interactions. However, the site-specificity of prolyl 4-hydroxylase 1 (P4ha1) and P4ha2 is not comprehensively studied yet. Further, the effect of P4ha1 and P4ha2 on the plethora of other site-specific collagen PTMs is not known to date. In-depth mass-spectrometry data (PXD008802) analysis of mice skin collagen I extracted from wild-type and different deletion mutants of C-P4hs revealed that partial or full deletion of prolyl 4-hydroxylases (P4ha1 and P4ha2) significantly decreases collagen deposition in ECM hinting towards perturbed biosynthesis. A total of ***421*** site-specific PTMs on fibrillar collagen chains (Col1a1, Col1a2, and Col3a1) were identified. Further, novel ***23*** P4ha1 specific, ***8*** P4ha2 specific, and ***18*** C-P4hs promiscuous sites on fibrillar collagen chains were identified. Partial deletion of P4ha1 and full deletion of P4ha2 also resulted in altered levels of the site-specific prolyl-3-hydroxylation occupancy in collagen I. Surprisingly, an increased level of site-specific lysyl hydroxylation (Col1a1-K^731^, Col1a2-K^183,315^) was documented upon partial deletion of P4ha1 and full deletion of P4ha2. Our findings showcased that the activity of prolyl 4-hydroxylases is not limited to 4-hydroxylation of specific proline sites, but simultaneously can perturb the entire biosynthetic network by modulating prolyl 3-hydroxylation and lysyl hydroxylation occupancy levels in the fibrillar collagen chains in a site-specific manner.

## Introduction

Collagens are the most abundant component of the extracellular matrix (ECM). Collagens provide an attachment surface to the cells in the body. Collagens maintain the structural stability and elasticity of all tissues [1–3]. Collagens play important roles in cellular functions like cell-to-cell communication, cell migration, and cell growth by interaction with cell surface receptors (integrins, discoidin domain receptor, glycoprotein VI, and FC gamma receptor) [4–7]. Collagens are very dynamic in structure. Interestingly, the complexity of the expression of collagen is not just limited to the differential abundance in a tissue-specific manner. Collagen complexity is extended to hundreds of tissue-specific post-translational modifications (PTMs) of collagen chains having tissue-specific variations. Collagens are heavily decorated with PTMs during biosynthesis in the endoplasmic reticulum (ER) prior to helix formation. PTMs are required for proper folding, stability, and functioning of the collagens [8–18]. Collagen PTMs are site-specifically catalyzed by collagen-modifying enzyme families. Collagen prolyl 4-hydroxylases (C-P4hs) i.e., Prolyl 4-hydroxylase alpha 1 (P4HA1), P4HA2, and P4HA3 catalyze 4-hydroxylation on prolines present in collagen chains. The 3-hydroxylation on proline is further catalyzed by prolyl 3-hydroxylase 1 (P3H1), P3H2, and P3H3 in the collagen chains. Additionally, lysines present in the collagens can also be 5-hydroxylated. Hydroxylysines in collagens also serve as a substrate for O-glycosylating enzymes. Lysyl hydroxylation in the collagen is catalyzed by lysyl hydroxylase 1 (LH1), LH2, and LH3 encoded by genes procollagen-lysine,2-oxoglutarate 5-dioxygenases (PLODs). O-glycosylation on hydroxylysines is catalyzed by procollagen galactosyltransferases (COLGALTs). Further, these galactosyl-hydroxylysine residues can also be modified with another glucose molecule via an o-glycosidic bond formation by glucosyltransferases. For the last five decades, several researchers have been putting effort to understand the detailed physiological role of these collagen PTMs [11,12,14–17,19–23]. However, the knowledge is still limited. One of the most well-studied collagen PTM is prolyl-4-hydroxylation [11,23–27]. The role of prolyl 4-hydroxylases have been studied in thermal stability and also in adverse ECM remodeling leading to pathophysiological complications such as fibrosis and cancer progression [28–32]. P4ha1 is the predominant isoform of collagen prolyl 4-hydroxylases. It has systemic expression in the body [23,33]. P4ha2 has higher expression in epithelial cells and bone tissues, and it is found to be elevated in many cancerous tissues [23,33]. P4ha3 is the least abundant isoform of collagen prolyl 4-hydroxylases [23]. It has been reported that complete deletion of P4ha1 (P4ha1−/−) is embryonically lethal [23,33]. However, mice with partial deletion of P4ha1 (P4ha1+/−) are viable. Mice with partial (P4ha2+/−) or complete (P4ha2−/−) deletion of P4ha2 are also viable with no apparent phenotype [23,33].

However, the comprehensive site-specificity of P4ha1 and P4ha2 is yet to be fully explored. Further, the effects of P4ha1 and P4ha2 deletion upon site-specific prolyl-3-hydroxylation, lysyl-hydroxylation, and O-glycosylation on collagen molecules are not known yet. So, here we utilized our *in-house* proteomics pipeline to study the effects of P4ha1 and P4ha2 enzymes on site-specific modifications and biosynthesis of fibrillar collagen chains (Col1a1, Col1a2, and COL3a1) extracted from the mice skin of different deletion mutants of C-P4hs. We utilized raw mass-spectrometry data of mice skin fibrillar collagen extracted from different deletion mutants of P4ha1 and P4ha2 (P4ha1+/−; P4ha2−/−, P4ha1+/+; P4ha2−/−, P4ha1+/−; P4ha2+/−, P4ha1+/+; P4ha2+/−, and wild type mice) to delineate the substrate specificity and overall effect on other collagen PTMs. Here we showcase that, P4ha1 and P4ha2 have site-specificity in the fibrillar collagens, and we have also shown for the first time that the effect of deletion of prolyl 4-hydroxylases is not just limited to the loss of 4-hydroxylation on specific proline sites, but it is also extended to affect to the entire fibrillar collagen PTM network.

## Methods

### Mass spectrometry data resource-

In this study, publicly available proteomic dataset #PXD008802 submitted by Sipilä *et. al.* 2018 [23] in ProteomeXchange was utilized. This dataset contains raw mass spectrometry files of fibrillar collagen extracted from the skin of prolyl 4-hydroxylase mutants and wild-type mice. There are 12 raw mass spectrometry files present for P4ha1+/−; P4ha2−/− group (n=6), 14 raw files for P4ha1+/+; P4ha2−/− (n=7), 10 raw files for P4ha1+/−; P4ha2+/− (n=5), 8 raw files for P4ha1+/+ (n=4); P4ha2+/−, and 10 raw files for wild type (n=5) in the #PXD008802 dataset. All the biological replicates had a technical duplicate. This proteomic dataset acquisition was performed using nanoflow HPLC system (Easy-nLC1000, Thermo Fisher Scientific) coupled with Q Exactive mass spectrometer from ThermoFisher as described earlier [23].

### Proteomic data analysis-

In this study, 54 raw mass spectrometry files from dataset #PXD008802 were analyzed using our *in-house* pipeline [13,34,35]. Two different search engines Andromeda (embedded within MaxQuant) and MyriMatch were employed for this analysis. We used MaxQuant for the relative quantification of collagen chains in wild-type and different prolyl 4-hydroxylase deletion mutants. MyriMatch was utilized to probe the site-specific identification of collagen PTMs (hydroxylation of prolines and lysines, O-glycosylation of hydroxylysines) in fibrillar collagen chains.

### Database search for the identification of collagens and site-specific collagen PTMs (Hydroxylation of proline and lysine, O-Glycosylation of lysines) using MyriMatch-

MyriMatch[36] was used for the identification of site-specific collagen PTMs. Firstly, a general database search was performed on *Mus musculus* database (.FASTA) downloaded from Uniprot.org (downloaded on 4-Dec-2021) containing 17090 entries. In general search, tolerance for precursor was set at ±10 ppm and for fragment ions, it was allowed up to ±20 ppm. Fully tryptic peptides were considered for database search with missed cleavages allowed up to a maximum of 2 per peptide. Carbamidomethylation (+57.0236) on cysteine was used as static modification and methionine oxidation (+15.994916) along with hydroxyproline (+15.994916) were used as dynamic modifications. Maximum dynamic modifications per peptide were set to a maximum of 4 per peptide. IDPICKER[37] was used for parsimonious grouping of the MyriMatch (*.pepXML) output file. FDR was controlled at <1% at PSMs, peptide, and protein group levels. After a general database search, the list of identified proteins was further exported from IDPicker as subset fasta. This subset *.FASTA database was used for further in-depth collagen PTM search using MyriMatch. In the subset FASTA database search using MyriMatch precursor and fragment ion mass tolerance was kept at 10 ppm and 20 ppm respectively as mentioned earlier [13,34,38]. Carbamidomethylation (+57.0236) of cysteine was used as static modification and Oxidation (+15.994916) of methionine was used as dynamic modification. Oxidation (+15.994916) of proline was used as a dynamic modification for the detection of hydroxyprolines in the full-length collagen chains. Further motif-based following dynamic modifications were also included; 3-prolyl-hydroxylation (GP!P! 15.994916), lysyl-hydroxylation (GXK! 15.994916), galactosyl-hydroxylysine (GXK! 178.047738) and glucosyl galactosyl-hydroxylysine (GXK! 340.100562). A maximum of 10 modifications were allowed per peptide. A maximum number of 4 missed cleavages were allowed for fully tryptic peptides. MyriMatch searches were completed to generate respective pepXML files. These pepXML files were further grouped for PSM matches, peptide, and protein group identification using IDPicker with <1% FDR (for PSM, peptide, and protein IDs). Identification of 3-hydroxyprolines was only considered if a proline residue was found to be hydroxylated at the “X” position of a “-Xaa-HyP-Gly” motif in the collagen chains. Further, pLabel [39] was used for manual inspection, analysis, and validation of subset database search PSMs for assigning a specific collagen PTM-containing peptide.

### Relative quantitation of abundances of collagen chains in wild-type and prolyl 4-hydroxylases mutant mice-

Quantitation of collagen chains from raw mass spectrometry data was performed with MaxQuant_ 2.2.0.0 [40]. Previously exported subset FASTA database was used for the searches conducted through MaxQuant (v 2.0.1.0). Up to 2 missed cleavages were allowed for the trypsin digestion. Carbamidomethylation (+57.0236) of cysteine was used as fixed modification and oxidation (+15.994916) of methionine, lysine, and proline along with N-terminal acetylation (42.010565) was used as variable modifications. Maximum modifications were set at 5 per peptide. The false discovery rate (FDR) was set at 0.01 for peptide and protein levels. The matches between the runs feature and LFQ module for relative quantitation were enabled. LFQ intensities of the three fibrillar collagen chains across different samples were computed and further used for relative quantitation.

### Quantitation of occupancy level of collagen PTM sites using Skyline-

MyriMatch search results (*.pepXML) were parsed through Peptide Prophet (TPP pipeline module)[41] for probability scoring between 0 to 1. Parsed MyriMatch results were utilized for building the spectral library in the Skyline[42]. The spectral library was utilized for the extraction of MS^1^ intensities of different unmodified and modified forms of collagen peptides from the raw mass spectrometry data as described previously [13,34,35].

### Statistical Analysis-

For the statistical analysis of the level of collagen chains and occupancy level of site-specific PTMs, a two-tailed Student’s test was applied. For correlation analysis, the Spearman test was conducted. A p-value of <0.05 was considered be to statistically significant in all tests.

## Results

Prolyl 4-hydroxylation is essential for the biosynthesis of collagens. It provides thermal stability to the collagen triple helix. P4HA1 is the predominant isoform catalyzing prolyl 4-hydroxylation on collagen chains. P4ha2 isoform also plays a critical role in collagen biosynthesis. To assess the impact of P4ha1 and P4ha2 deletion on the deposition of collagen in the ECM, we quantitated the level of fibrillar collagen chains (COL1A1, COL1A2, and COL3A1) from the skin of wild-type and different deletion mutants of C-P4hs.

### 3.1 Relative abundance of fibrillar collagen chains from the skin of wild-type and different C-P4hs deletion mutant mice-

The relative label-free quantitation of collagen 1 chains (Col1a1 and Col1a2) revealed that deletion of prolyl 4-hydroxylases have adversely affected the collagen deposition in the ECM of skin. Previously Tolonen et. al., [33] have found significant decrease in collagen amount in mice bone upon complete deletion of P4ha2. Here in our analysis of mice skin, we found lower levels of collagen 1 in prolyl 4-hydroxylases deleted mice. We detected a significant (p<0.05) ∼10% decrease in the Col1a1 level in mice having a partial deletion of P4ha2 (P4ha1+/+; P4ha2+/−) and mice having a partial deletion of P4ha1 and P4ha2 (P4HA1+/−; P4HA2+/−) compared to wild type mice. However, we detected ∼15.5% significant (p<0.05) decrease in Col1a1 level in mice having complete deletion of P4ha2 (P4ha1+/+; P4ha2−/−). Col1a1 level was also significantly (p<0.05) decreased by about 17% in mice having a partial deletion of P4ha1 and complete deletion of P4ha2 (P4ha1+/−; P4ha2−/−) compared to the wild-type mice (Figure 2A). Similarly, the level of Col1a2 was also significantly (p<0.05) decreased by ∼8.5% in P4ha1+/+; P4ha2+/− mice compared to wild-type. Col1a2 was also found to be significantly (p<0.05) decreased ∼11% in P4ha1+/−; P4ha2+/− mice compared to wild-type. Col1a2 level in skin ECM was further significantly decreased (p<0.05) ∼18.5% in P4ha1+/+; P4ha2−/− mice and it decreased ∼23% in P4ha1+/−; P4ha2−/− mice compared to wild type mice (Figure 2B). We found that complete deletion of P4ha2 resulted in lower collagen 1 levels compared to wild type or partial deletion of P4ha1 or partial deletion of P4ha2. P4ha1+/−; P4ha2−/− mice were found to have the lowest amount of collagen 1 chains deposited in the skin thereby implicating a potential decrease in collagen 1 biosynthesis. Although collagen 1 was less deposited in P4ha1+/−; P4ha2−/− mice, we detected a significant (p<0.05) elevation of about 34% in the level of Col3a1 in P4ha1+/−; P4ha2−/− mice compared to the wild-type (Figure 2C).

**Figure 1:**
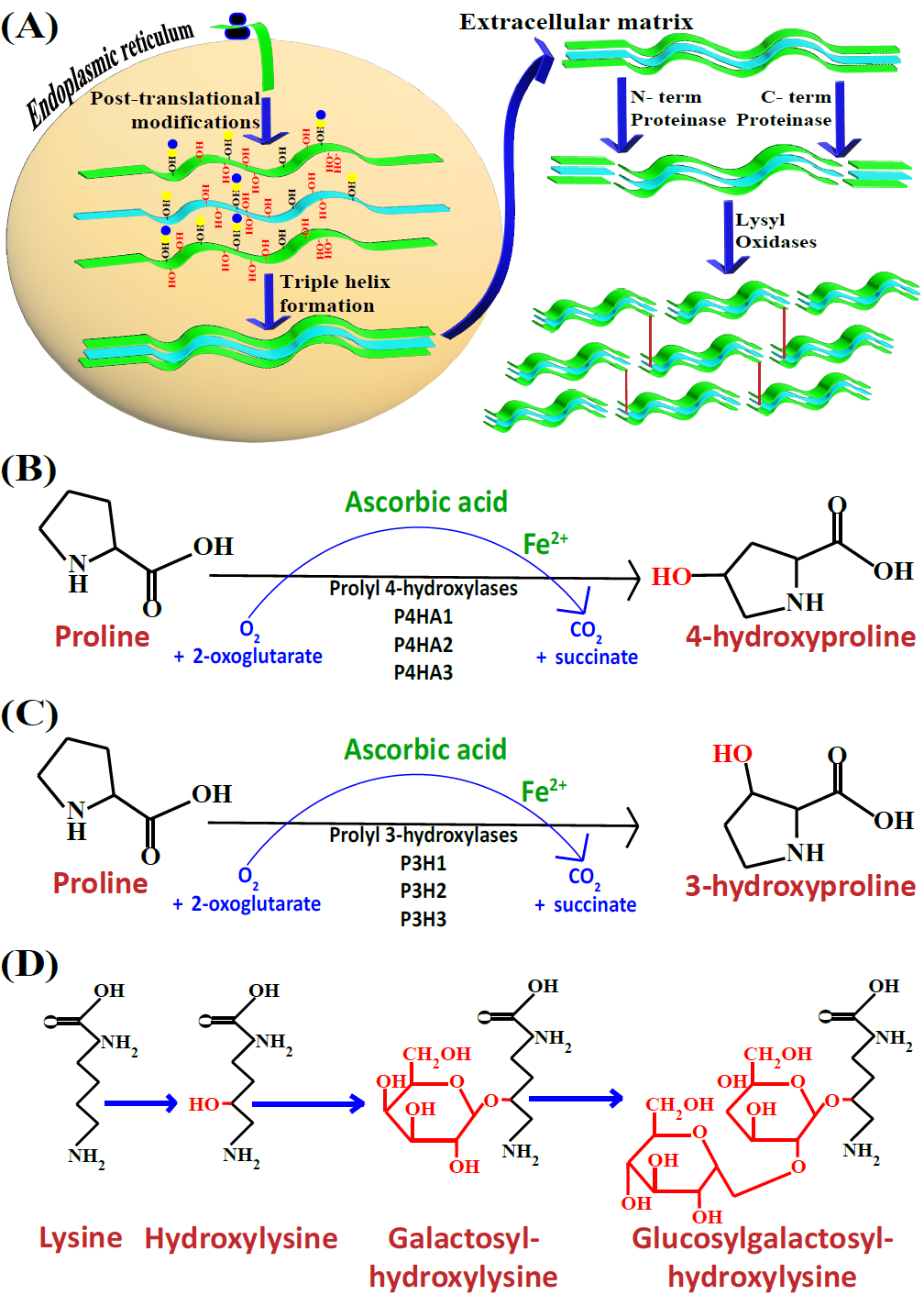
Occurrence of collagen PTMs during biosynthesis and deposition in the ECM-. Figure 1(A) shows the biosynthesis and deposition of collagen in the ECM. Newly translated collagen chains enter the endoplasmic reticulum and get heavily modified post-translationally in a site-specific manner by collagen-modifying enzymes. Modified collagen chains form a triple helix, which is transported to the ECM. In ECM, N and C terminal proteinases cleave the propeptides of the collagen triple helix. Then cross-linking and formation of collagen fibre assembly is induced by the activity of lysyl-oxidases. Figure 1(B) shows the formation of 4-hydroxyproline from proline catalyzed by prolyl 4-hydroxylases in the presence of oxygen and 2-oxoglutarate with ascorbic acid and ferric cation co-factors. Figure 1(C), the process of formation of 3-hydroxyproline is similar to the formation of 4-hydroxyproline, however, prolyl 3-hydroxylases have different motif specificity than the prolyl 4-hydroxylases. Figure 1(D), with a process similar to prolyl 4-hydroxylases and prolyl 3-hydroxylases, lysyl hydroxylases can also 5-hydroxylate lysine present in collagen chains. 5-hydroxylysine serves as a substrate for O-glycosylation with galactosyl or glucosylgalactosyl by galactosyl and glucosyl transferases.

**Figure 2:**
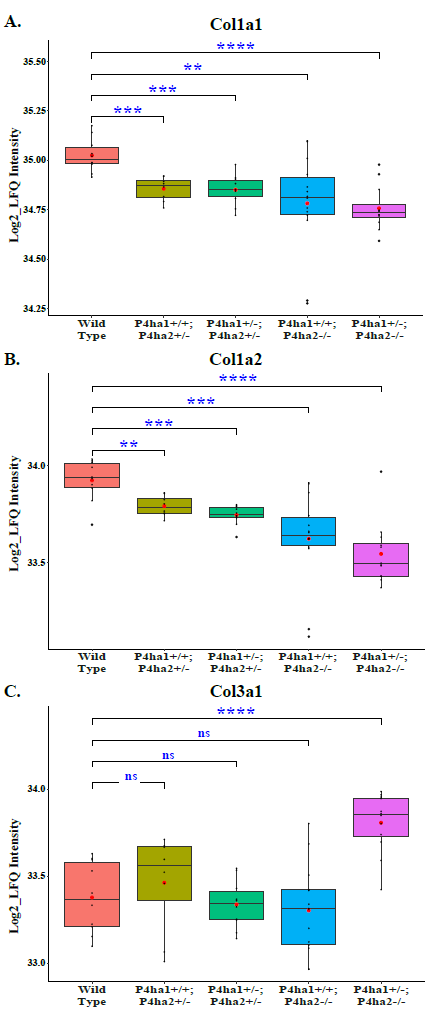
Relative abundance of fibrillar collagen chains deposited in mice skin ECM across different deletion mutants of C-P4hs and wild type-. (A and B) Partial deletion of P4ha1 and partial or complete deletion of P4ha2 reduced the collagen 1 (Col1a1 and Col1a2) deposition in the ECM. (C) Col3a1 was significantly elevated in P4ha1+/−; P4ha2−/− mice compared to the wild-types.

### 3.2 Identification of site-specific collagen PTMs in fibrillar collagens extracted from wild-type mice skin-

Fibrillar collagens, mainly collagen 1, have a high abundance in the ECM. Collagen I have the highest amount of hydroxyproline residues [34]. We identified hydroxyproline (HyP), 4-hydroxyproline (4-HyP), 3-hydroxyproline (3-HyP), 5-hydroxylysine (HyK), galactosyl-hydroxylysine (G-HyK) and glucosyl galactosyl-hydroxylysine (GG-HyK) sites on Col1a1, Col1a2 and Col3a1 chains from the skin of wild type mice (Table 1). Hydroxyproline present on the Yaa position of the -Xaa-Yaa-Gly motif was considered as 4-hydroxyproline. Hydroxyproline present on the Xaa position of -Xaa-4HyP-Gly-motif was considered as 3-hydroxyproline. Hydroxyproline present on Xaa position of -Xaa-Yaa-Gly-motif where Yaa is not a hydroxyproline/proline residue, are labeled as only “hydroxyproline” sites in this study. However, these sites are also likely to be 4-hydroxyproline in nature [43]. Hydroxylysine present on Yaa position of -Xaa-Yaa-Gly was considered as 5-hydroxylysine and these 5-hydroxylysines were also assessed for the presence of O-glycosylation modifications. On wild-type mice skin Col1a1, we detected a total of 160 site-specific PTMs. We identified a total of 106 4-hydroxyproline sites, 12 3-hydroxyproline sites, 22 hydroxyproline sites, 14 5-hydroxylysine sites, 4 galactosyl-hydroxylysine sites, and 2 glucosyl galactosyl-hydroxylysine sites on Col1a1.Col1a2 extracted from wild-type mice skin, revealed a total of 124 PTM sites in our analysis. A total of 89 4-hydroxyproline sites, 4 3-hydroxyproline sites, 17 hydroxyproline sites, and 14 5-hydroxylysine sites were detected on Col1a2. No O-glycosylation site on Col1a2 was detected from the skin of wild-type mice. Except for these two chains of collagen I, we also mapped PTMs on collagen III which forms a homotrimer of Col3a1 chains. We identified a total of 137 PTMs on wild-type mice skin Col3a1. We detected 112 4-hydroxyproline sites, 2 3-hydroxyproline sites, 10 hydroxyproline sites, 12 5-hydroxylysine sites, and 1 glucosyl-galactosyl-hydroxylysine sites on Col3a1. In our analysis, we identified a total of 421 site-specific collagen PTMs on Col1a1, Col1a2, and Col3a1 chains extracted from wild-type mice skin. We found Col1a1 to be the most modified and Col1a2 to be the least modified among the three fibrillar collagen chains.

**Table 1:**
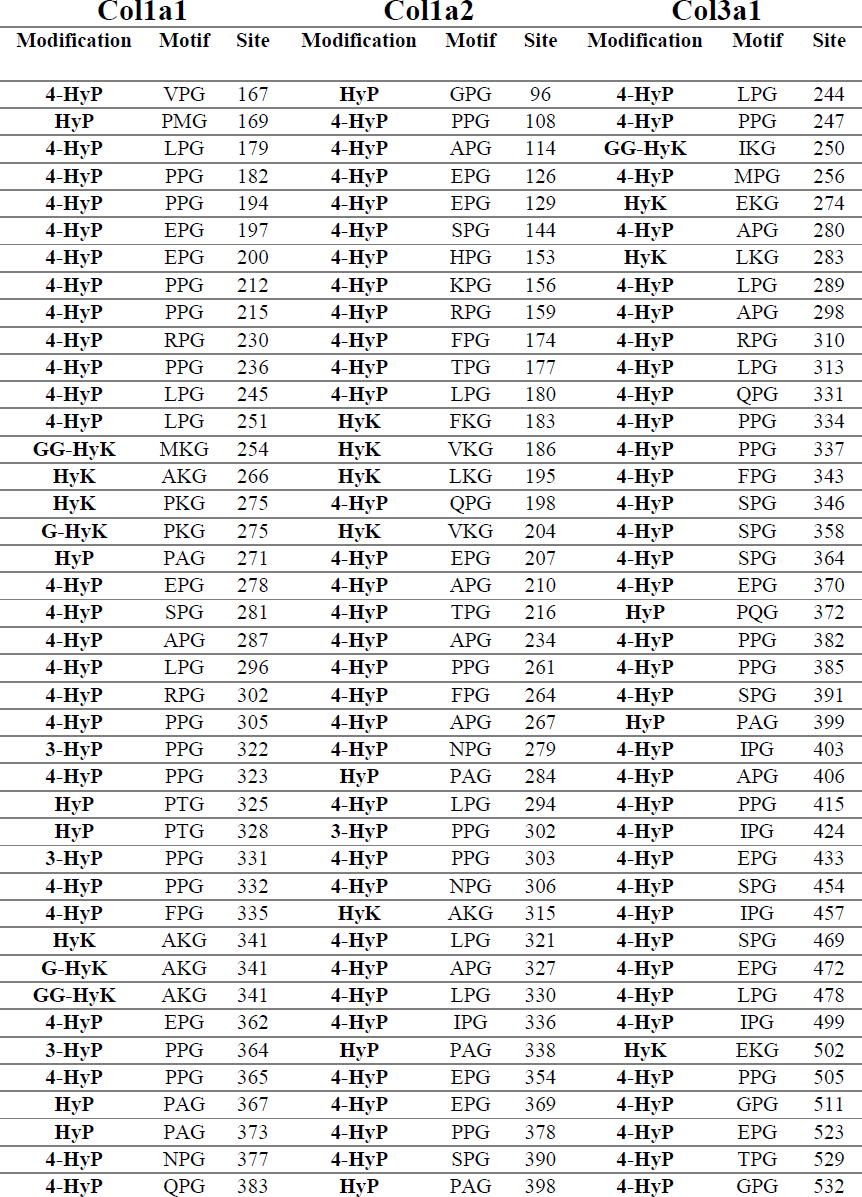
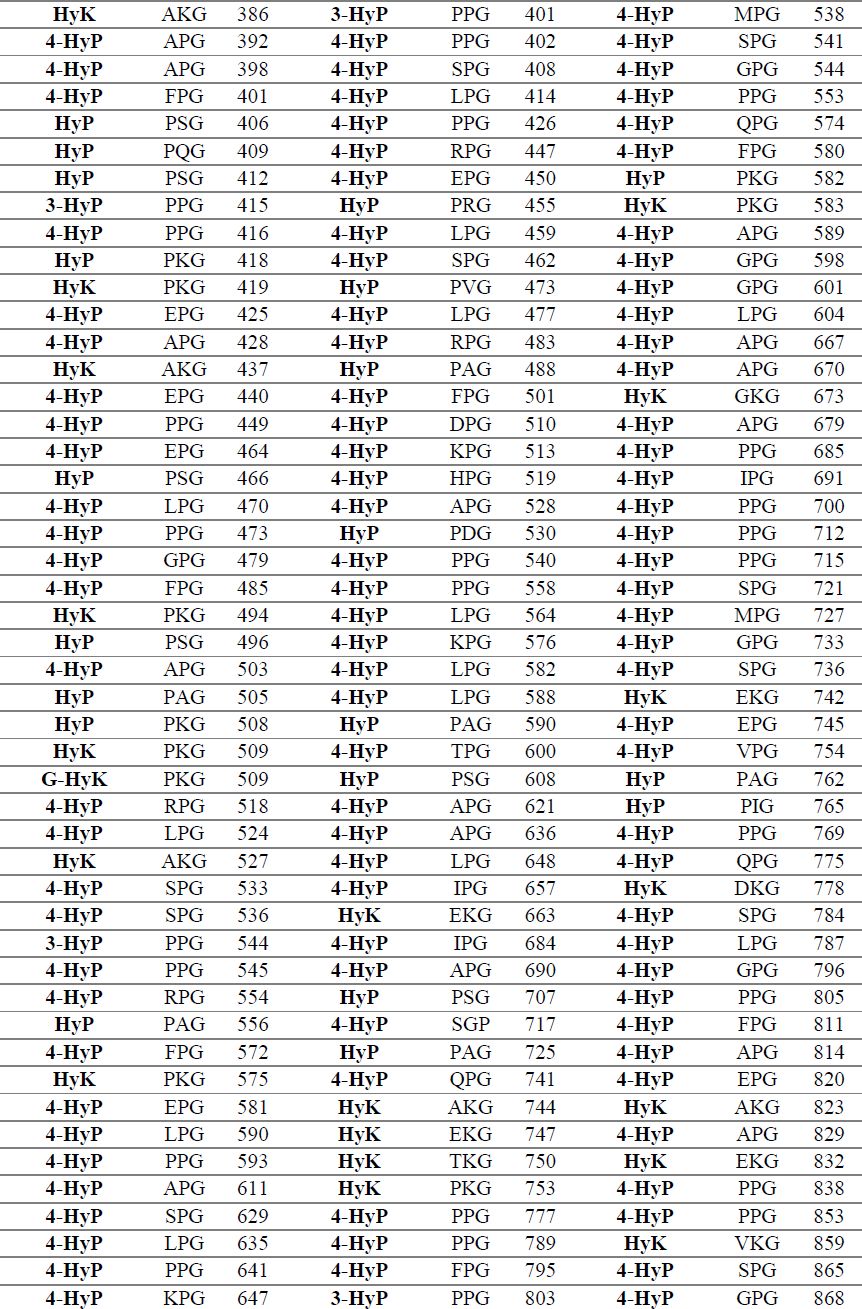
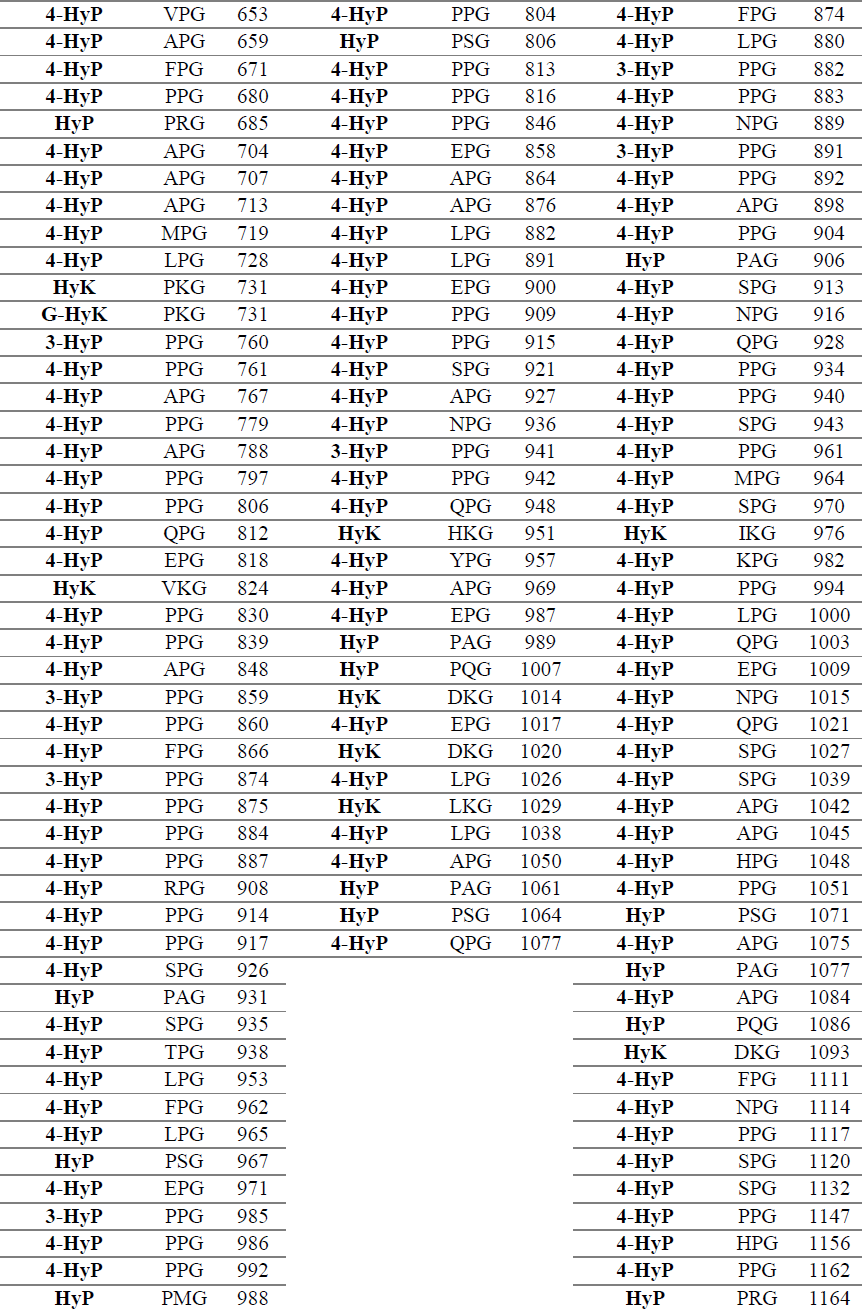
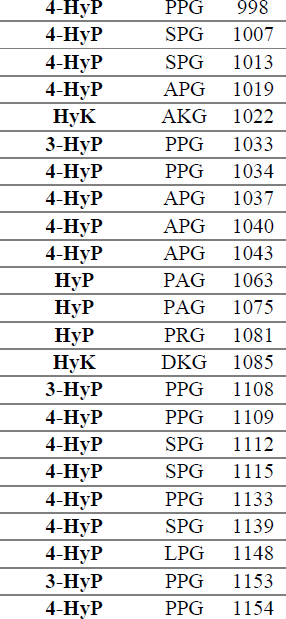
Site-specific collagen PTMs identified on fibrillar collagen chains extracted from wild-type mice skin.

| **Colla1** |  |  | **Colla2** |  |  | **Col3a1** |  |  |
| --- | --- | --- | --- | --- | --- | --- | --- | --- |
| **Modification** | **Motif** | **Site** | **Modification** | **Motif** | **Site** | **Modification** | **Motif** | **Site** |
| **4-HyP** | VPG | 167 | **HyP** | GPG | 96 | **4-HyP** | LPG | 244 |
| **HyP** | PMG | 169 | **4-HyP** | PPG | 108 | **4-HyP** | PPG | 247 |
| **4-HyP** | LPG | 179 | **4-HyP** | APG | 114 | **GG-HyK** | IKG | 250 |
| **4-HyP** | PPG | 182 | **4-HyP** | EPG | 126 | **4-HyP** | MPG | 256 |
| **4-HyP** | PPG | 194 | **4-HyP** | EPG | 129 | **HyK** | EKG | 274 |
| **4-HyP** | EPG | 197 | **4-HyP** | SPG | 144 | **4-HyP** | APG | 280 |
| **4-HyP** | EPG | 200 | **4-HyP** | HPG | 153 | **HyK** | LKG | 283 |
| **4-HyP** | PPG | 212 | **4-HyP** | KPG | 156 | **4-HyP** | LPG | 289 |
| **4-HyP** | PPG | 215 | **4-HyP** | RPG | 159 | **4-HyP** | APG | 298 |
| **4-HyP** | RPG | 230 | **4-HyP** | FPG | 174 | **4-HyP** | RPG | 310 |
| **4-HyP** | PPG | 236 | **4-HyP** | TPG | 177 | **4-HyP** | LPG | 313 |
| **4-HyP** | LPG | 245 | **4-HyP** | LPG | 180 | **4-HyP** | QPG | 331 |
| **4-HyP** | LPG | 251 | **HyK** | FKG | 183 | **4-HyP** | PPG | 334 |
| **GG-HyK** | MKG | 254 | **HyK** | VKG | 186 | **4-HyP** | PPG | 337 |
| **HyK** | AKG | 266 | **HyK** | LKG | 195 | **4-HyP** | FPG | 343 |
| **HyK** | PKG | 275 | **4-HyP** | QPG | 198 | **4-HyP** | SPG | 346 |
| **G-HyK** | PKG | 275 | **HyK** | VKG | 204 | **4-HyP** | SPG | 358 |
| **HyP** | PAG | 271 | **4-HyP** | EPG | 207 | **4-HyP** | SPG | 364 |
| **4-HyP** | EPG | 278 | **4-HyP** | APG | 210 | **4-HyP** | EPG | 370 |
| **4-HyP** | SPG | 281 | **4-HyP** | TPG | 216 | **HyP** | PQG | 372 |
| **4-HyP** | APG | 287 | **4-HyP** | APG | 234 | **4-HyP** | PPG | 382 |
| **4-HyP** | LPG | 296 | **4-HyP** | PPG | 261 | **4-HyP** | PPG | 385 |
| **4-HyP** | RPG | 302 | **4-HyP** | FPG | 264 | **4-HyP** | SPG | 391 |
| **4-HyP** | PPG | 305 | **4-HyP** | APG | 267 | **HyP** | PAG | 399 |
| **3-HyP** | PPG | 322 | **4-HyP** | NPG | 279 | **4-HyP** | IPG | 403 |
| **4-HyP** | PPG | 323 | **HyP** | PAG | 284 | **4-HyP** | APG | 406 |
| **HyP** | PTG | 325 | **4-HyP** | LPG | 294 | **4-HyP** | PPG | 415 |
| **HyP** | PTG | 328 | **3-HyP** | PPG | 302 | **4-HyP** | IPG | 424 |
| **3-HyP** | PPG | 331 | **4-HyP** | PPG | 303 | **4-HyP** | EPG | 433 |
| **4-HyP** | PPG | 332 | **4-HyP** | NPG | 306 | **4-HyP** | SPG | 454 |
| **4-HyP** | FPG | 335 | **HyK** | AKG | 315 | **4-HyP** | IPG | 457 |
| **HyK** | AKG | 341 | **4-HyP** | LPG | 321 | **4-HyP** | SPG | 469 |
| **G-HyK** | AKG | 341 | **4-HyP** | APG | 327 | **4-HyP** | EPG | 472 |
| **GG-HyK** | AKG | 341 | **4-HyP** | LPG | 330 | **4-HyP** | LPG | 478 |
| **4-HyP** | EPG | 362 | **4-HyP** | IPG | 336 | **4-HyP** | IPG | 499 |
| **3-HyP** | PPG | 364 | **HyP** | PAG | 338 | **HyK** | EKG | 502 |
| **4-HyP** | PPG | 365 | **4-HyP** | EPG | 354 | **4-HyP** | PPG | 505 |
| **HyP** | PAG | 367 | **4-HyP** | EPG | 369 | **4-HyP** | GPG | 511 |
| **HyP** | PAG | 373 | **4-HyP** | PPG | 378 | **4-HyP** | EPG | 523 |
| **4-HyP** | NPG | 377 | **4-HyP** | SPG | 390 | **4-HyP** | TPG | 529 |
| **4-HyP** | QPG | 383 | **HyP** | PAG | 398 | **4-HyP** | GPG | 532 |
| **HyK** | AKG | 386 | **3-HyP** | PPG | 401 | **4-HyP** | MPG | 538 |
| **4-HyP** | APG | 392 | **4-HyP** | PPG | 402 | **4-HyP** | SPG | 541 |
| **4-HyP** | APG | 398 | **4-HyP** | SPG | 408 | **4-HyP** | GPG | 544 |
| **4-HyP** | FPG | 401 | **4-HyP** | LPG | 414 | **4-HyP** | PPG | 553 |
| **HyP** | PSG | 406 | **4-HyP** | PPG | 426 | **4-HyP** | QPG | 574 |
| **HyP** | PQG | 409 | **4-HyP** | RPG | 447 | **4-HyP** | FPG | 580 |
| **HyP** | PSG | 412 | **4-HyP** | EPG | 450 | **HyP** | PKG | 582 |
| **3-HyP** | PPG | 415 | **HyP** | PRG | 455 | **HyK** | PKG | 583 |
| **4-HyP** | PPG | 416 | **4-HyP** | LPG | 459 | **4-HyP** | APG | 589 |
| **HyP** | PKG | 418 | **4-HyP** | SPG | 462 | **4-HyP** | GPG | 598 |
| **HyK** | PKG | 419 | **HyP** | PVG | 473 | **4-HyP** | GPG | 601 |
| **4-HyP** | EPG | 425 | **4-HyP** | LPG | 477 | **4-HyP** | LPG | 604 |
| **4-HyP** | APG | 428 | **4-HyP** | RPG | 483 | **4-HyP** | APG | 667 |
| **HyK** | AKG | 437 | **HyP** | PAG | 488 | **4-HyP** | APG | 670 |
| **4-HyP** | EPG | 440 | **4-HyP** | FPG | 501 | **HyK** | GKG | 673 |
| **4-HyP** | PPG | 449 | **4-HyP** | DPG | 510 | **4-HyP** | APG | 679 |
| **4-HyP** | EPG | 464 | **4-HyP** | KPG | 513 | **4-HyP** | PPG | 685 |
| **HyP** | PSG | 466 | **4-HyP** | HPG | 519 | **4-HyP** | IPG | 691 |
| **4-HyP** | LPG | 470 | **4-HyP** | APG | 528 | **4-HyP** | PPG | 700 |
| **4-HyP** | PPG | 473 | **HyP** | PDG | 530 | **4-HyP** | PPG | 712 |
| **4-HyP** | GPG | 479 | **4-HyP** | PPG | 540 | **4-HyP** | PPG | 715 |
| **4-HyP** | FPG | 485 | **4-HyP** | PPG | 558 | **4-HyP** | SPG | 721 |
| **HyK** | PKG | 494 | **4-HyP** | LPG | 564 | **4-HyP** | MPG | 727 |
| **HyP** | PSG | 496 | **4-HyP** | KPG | 576 | **4-HyP** | GPG | 733 |
| **4-HyP** | APG | 503 | **4-HyP** | LPG | 582 | **4-HyP** | SPG | 736 |
| **HyP** | PAG | 505 | **4-HyP** | LPG | 588 | **HyK** | EKG | 742 |
| **HyP** | PKG | 508 | **HyP** | PAG | 590 | **4-HyP** | EPG | 745 |
| **HyK** | PKG | 509 | **4-HyP** | TPG | 600 | **4-HyP** | VPG | 754 |
| **G-HyK** | PKG | 509 | **HyP** | PSG | 608 | **HyP** | PAG | 762 |
| **4-HyP** | RPG | 518 | **4-HyP** | APG | 621 | **HyP** | PIG | 765 |
| **4-HyP** | LPG | 524 | **4-HyP** | APG | 636 | **4-HyP** | PPG | 769 |
| **HyK** | AKG | 527 | **4-HyP** | LPG | 648 | **4-HyP** | QPG | 775 |
| **4-HyP** | SPG | 533 | **4-HyP** | IPG | 657 | **HyK** | DKG | 778 |
| **4-HyP** | SPG | 536 | **HyK** | EKG | 663 | **4-HyP** | SPG | 784 |
| **3-HyP** | PPG | 544 | **4-HyP** | IPG | 684 | **4-HyP** | LPG | 787 |
| **4-HyP** | PPG | 545 | **4-HyP** | APG | 690 | **4-HyP** | GPG | 796 |
| **4-HyP** | RPG | 554 | **HyP** | PSG | 707 | **4-HyP** | PPG | 805 |
| **HyP** | PAG | 556 | **4-HyP** | SGP | 717 | **4-HyP** | FPG | 811 |
| **4-HyP** | FPG | 572 | **HyP** | PAG | 725 | **4-HyP** | APG | 814 |
| **HyK** | PKG | 575 | **4-HyP** | QPG | 741 | **4-HyP** | EPG | 820 |
| **4-HyP** | EPG | 581 | **HyK** | AKG | 744 | **HyK** | AKG | 823 |
| **4-HyP** | LPG | 590 | **HyK** | EKG | 747 | **4-HyP** | APG | 829 |
| **4-HyP** | PPG | 593 | **HyK** | TKG | 750 | **HyK** | EKG | 832 |
| **4-HyP** | APG | 611 | **HyK** | PKG | 753 | **4-HyP** | PPG | 838 |
| **4-HyP** | SPG | 629 | **4-HyP** | PPG | 777 | **4-HyP** | PPG | 853 |
| **4-HyP** | LPG | 635 | **4-HyP** | PPG | 789 | **HyK** | VKG | 859 |
| **4-HyP** | PPG | 641 | **4-HyP** | FPG | 795 | **4-HyP** | SPG | 865 |
| **4-HyP** | KPG | 647 | **3-HyP** | PPG | 803 | **4-HyP** | GPG | 868 |
| **4-HyP** | VPG | 653 | **4-HyP** | PPG | 804 | **4-HyP** | FPG | 874 |
| **4-HyP** | APG | 659 | **HyP** | PSG | 806 | **4-HyP** | LPG | 880 |
| **4-HyP** | FPG | 671 | **4-HyP** | PPG | 813 | **3-HyP** | PPG | 882 |
| **4-HyP** | PPG | 680 | **4-HyP** | PPG | 816 | **4-HyP** | PPG | 883 |
| **HyP** | PRG | 685 | **4-HyP** | PPG | 846 | **4-HyP** | NPG | 889 |
| **4-HyP** | APG | 704 | **4-HyP** | EPG | 858 | **3-HyP** | PPG | 891 |
| **4-HyP** | APG | 707 | **4-HyP** | APG | 864 | **4-HyP** | PPG | 892 |
| **4-HyP** | APG | 713 | **4-HyP** | APG | 876 | **4-HyP** | APG | 898 |
| **4-HyP** | MPG | 719 | **4-HyP** | LPG | 882 | **4-HyP** | PPG | 904 |
| **4-HyP** | LPG | 728 | **4-HyP** | LPG | 891 | **HyP** | PAG | 906 |
| **HyK** | PKG | 731 | **4-HyP** | EPG | 900 | **4-HyP** | SPG | 913 |
| **G-HyK** | PKG | 731 | **4-HyP** | PPG | 909 | **4-HyP** | NPG | 916 |
| **3-HyP** | PPG | 760 | **4-HyP** | PPG | 915 | **4-HyP** | QPG | 928 |
| **4-HyP** | PPG | 761 | **4-HyP** | SPG | 921 | **4-HyP** | PPG | 934 |
| **4-HyP** | APG | 767 | **4-HyP** | APG | 927 | **4-HyP** | PPG | 940 |
| **4-HyP** | PPG | 779 | **4-HyP** | NPG | 936 | **4-HyP** | SPG | 943 |
| **4-HyP** | APG | 788 | **3-HyP** | PPG | 941 | **4-HyP** | PPG | 961 |
| **4-HyP** | PPG | 797 | **4-HyP** | PPG | 942 | **4-HyP** | MPG | 964 |
| **4-HyP** | PPG | 806 | **4-HyP** | QPG | 948 | **4-HyP** | SPG | 970 |
| **4-HyP** | QPG | 812 | **HyK** | HKG | 951 | **HyK** | IKG | 976 |
| **4-HyP** | EPG | 818 | **4-HyP** | YPG | 957 | **4-HyP** | KPG | 982 |
| **HyK** | VKG | 824 | **4-HyP** | APG | 969 | **4-HyP** | PPG | 994 |
| **4-HyP** | PPG | 830 | **4-HyP** | EPG | 987 | **4-HyP** | LPG | 1000 |
| **4-HyP** | PPG | 839 | **HyP** | PAG | 989 | **4-HyP** | QPG | 1003 |
| **4-HyP** | APG | 848 | **HyP** | PQG | 1007 | **4-HyP** | EPG | 1009 |
| **3-HyP** | PPG | 859 | **HyK** | DKG | 1014 | **4-HyP** | NPG | 1015 |
| **4-HyP** | PPG | 860 | **4-HyP** | EPG | 1017 | **4-HyP** | QPG | 1021 |
| **4-HyP** | FPG | 866 | **HyK** | DKG | 1020 | **4-HyP** | SPG | 1027 |
| **3-HyP** | PPG | 874 | **4-HyP** | LPG | 1026 | **4-HyP** | SPG | 1039 |
| **4-HyP** | PPG | 875 | **HyK** | LKG | 1029 | **4-HyP** | APG | 1042 |
| **4-HyP** | PPG | 884 | **4-HyP** | LPG | 1038 | **4-HyP** | APG | 1045 |
| **4-HyP** | PPG | 887 | **4-HyP** | APG | 1050 | **4-HyP** | HPG | 1048 |
| **4-HyP** | RPG | 908 | **HyP** | PAG | 1061 | **4-HyP** | PPG | 1051 |
| **4-HyP** | PPG | 914 | **HyP** | PSG | 1064 | **HyP** | PSG | 1071 |
| **4-HyP** | PPG | 917 | **4-HyP** | QPG | 1077 | **4-HyP** | APG | 1075 |
| **4-HyP** | SPG | 926 |  |  |  | **HyP** | PAG | 1077 |
| **HyP** | PAG | 931 |  |  |  | **4-HyP** | APG | 1084 |
| **4-HyP** | SPG | 935 |  |  |  | **HyP** | PQG | 1086 |
| **4-HyP** | TPG | 938 |  |  |  | **HyK** | DKG | 1093 |
| **4-HyP** | LPG | 953 |  |  |  | **4-HyP** | FPG | 1111 |
| **4-HyP** | FPG | 962 |  |  |  | **4-HyP** | NPG | 1114 |
| **4-HyP** | LPG | 965 |  |  |  | **4-HyP** | PPG | 1117 |
| **HyP** | PSG | 967 |  |  |  | **4-HyP** | SPG | 1120 |
| **4-HyP** | EPG | 971 |  |  |  | **4-HyP** | SPG | 1132 |
| **3-HyP** | PPG | 985 |  |  |  | **4-HyP** | PPG | 1147 |
| **4-HyP** | PPG | 986 |  |  |  | **4-HyP** | HPG | 1156 |
| **4-HyP** | PPG | 992 |  |  |  | **4-HyP** | PPG | 1162 |
| **HyP** | PMG | 988 |  |  |  | **HyP** | PRG | 1164 |

|  |  |  |
| --- | --- | --- |
| 4-HyP | PPG | 998 |
| 4-HyP | SPG | 1007 |
| 4-HyP | SPG | 1013 |
| 4-HyP | APG | 1019 |
| HyK | AKG | 1022 |
| 3-HyP | PPG | 1033 |
| 4-HyP | PPG | 1034 |
| 4-HyP | APG | 1037 |
| 4-HyP | APG | 1040 |
| 4-HyP | APG | 1043 |
| HyP | PAG | 1063 |
| HyP | PAG | 1075 |
| HyP | PRG | 1081 |
| HyK | DKG | 1085 |
| 3-HyP | PPG | 1108 |
| 4-HyP | PPG | 1109 |
| 4-HyP | SPG | 1112 |
| 4-HyP | SPG | 1115 |
| 4-HyP | PPG | 1133 |
| 4-HyP | SPG | 1139 |
| 4-HyP | LPG | 1148 |
| 3-HyP | PPG | 1153 |
| 4-HyP | PPG | 1154 |

### 3.3 Mass Spectrometry-based validation of -Xaa-Pro-Gly motif catalyzed by prolyl 4-hydroxylases-

There are -Gly-Xaa-Yaa-repeats present in the helical region of collagen 1. During the proteomics analyses hydroxyproline present on Yaa position of -Gly-Xaa-Yaa- or -Gly-Xaa-Yaa-Gly-motif is considered to be 4-hydroxylated. On wild-type mice skin Col1a1, out of **106** 4-hydroxyproline sites, **17** 4-hydroxyproline sites were detected on -Ala-Pro-Gly-motif, **10** 4-hydroxyproline sites are detected on -Glu-Pro-Gly-motif, **7** 4-hydroxyproline sites are detected on -Phe-Pro-Gly-motif. In our analysis, the most common motif for 4-hydroxyproline is the -Pro-Pro-Gly-motif. A total of **35** 4-hydroxyproline sites on this motif were identified. We detected **12** 4-hydroxyproline sites on the -Leu-Pro-Gly-motif, **5** 4-hydroxyproline sites on the -Arg-Pro-Gly-motif, **11** 4-hydroxyproline sites on the -Ser-Pro-Gly-motif, **2-2** 4-hydroxyproline sites on the -Gln-Pro-Gly-motif and -Val-Pro-Gly-motifs. We also detected **1** site each on the -Gly-Pro-Gly-motif, -Lys-Pro-Gly-, -Met-Pro-Gly-, -Asn-Pro-Gly-motif, and -Thr-Pro-Gly-motif. Initial studies on the enzyme activity of C-P4h showed that the -Xaa-Yaa-Gly-motif is recognized by these enzymes and hydroxyproline present on the Yaa position of -Xaa-Yaa-Gly-motif is modified by the C-P4h [44–48]. Here, we validate these findings with the MS analysis. In col1a1, all 106 4-HyP sites indeed follow the repeat motif. We found hydroxyproline on the Yaa position of uniquely present -Xaa-Yaa-Gly-motif in both chains of collagen 1 at the starting of the helical region (Col1a1^161-176^, Col1a2^91-105^) confirming the Xaa-Yaa-Gly motif specificity of C4Phs as mentioned earlier. Col1a1 peptide (“^161^SAGVSVP_+16_GPMGPSGPR^176”^) having an m/z value of 734.8659^+2^ eluted at **9.21** minutes which is earlier than the unmodified version with an m/z value of 726.8654^+2^ (eluted at **9.60** minutes). Distinct y_10_ fragment ions of m/z values of 968.46^+1^ and 952.47^+1^ were the most abundant in the respective MS/MS spectra of the modified and unmodified versions of the peptide. Similarly, the Col1a2 peptide (“^91^GVSSGP_+16_GPMGLMGPR^105”^) having an m/z value of 708.3402^+2^ eluted at 10.78 minutes which is earlier than the unmodified version with an m/z value of 700.3448^+2^ (eluted at 11.08 minutes). Distinct y_10_ fragment ions of m/z values of 514.75^+2^, 1028.5^+1^ and 506.76^+2^, 1012.51^+^ were the most abundant in the respective MS/MS spectra of the modified and unmodified version of the peptide. On mice skin Col1a1 P167 we found hydroxyproline to be in -Xaa-HyP-Gly-motif (Figure 3 A2). Similarly, on mice skin Col1a2 P96, we found hydroxyproline to be also in -Xaa-HyP-Gly motif (Figure 3 B2). These -Xaa-Yaa-Gly-motifs are not part of -Gly-Xaa-Yaa- or -Gly-Xaa-Yaa-Gly-repeat motifs. So, the identification of these two hydroxyproline sites on -Xaa-Yaa-Gly-motif validates the initial finding on collagen prolyl 4-hydroxylase motif-specific activity.

**Figure 3:**
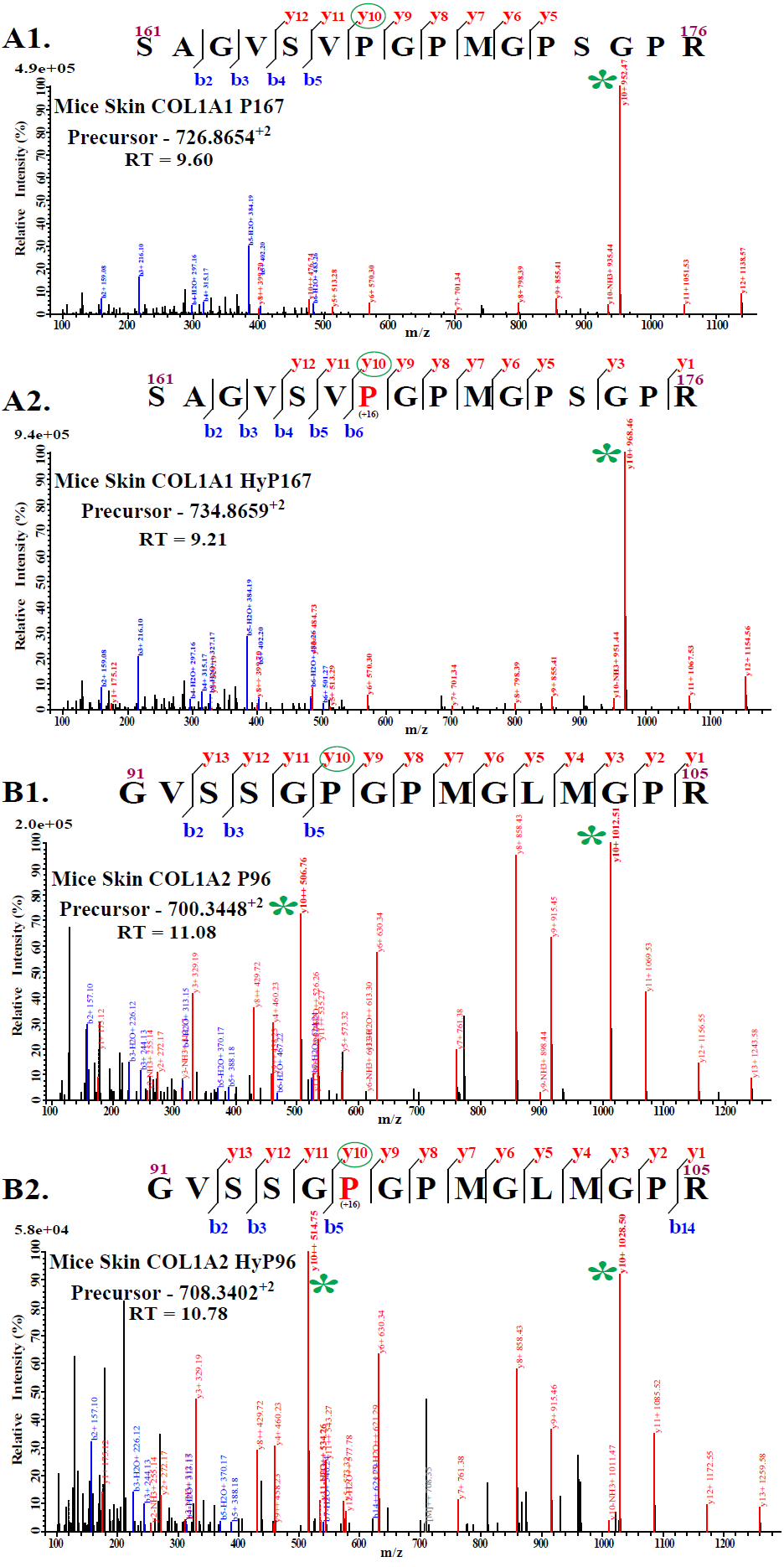
Validation of C-P4h specific motif (-Xaa-Pro-Gly-)-. A1. shows the unmodified peptide from Col1a1 having residues numbered 161-176. A2 shows the presence of hydroxyproline on P167, the presence of hydroxyproline is on the -Xaa-Pro-Gly-motif. Similarly, B1. Shows unmodified peptide from Col1a2 (91-105) and B2 shows the presence of hydroxyproline on P96 present on “Yaa” position of -Xaa-Yaa-Gly-motif.

### 3.4 Identification of site-specificity of C-P4hs in fibrillar collagens-

C-P4hs modify proline residues present on -Xaa-Pro-Gly-motif in collagens. However, collagen-modifying enzymes are known to have site-specificity [16,18,19,21,49–51]. The P3H1 knockout study revealed P3H1 has 3 specific sites in collagen I, these sites are majorly modified by P3H1. 3-hydroxyproline occupancy on these 3 sites is significantly diminished upon lack of P3H1 activity [18,49]. Similarly, P3H2 also has site-specificity in collagen IV [16,21]. However, the site-specificity of P4ha1 and P4ha2 in fibrillar collagens is yet to be fully explored. So, to identify P4ha1 and P4ha2 specific sites in mice skin fibrillar collagen chains, we quantitated the 4-HyP occupancy levels in wild-type and different C-P4h deletion mutants.

#### 3.4.1 Identification of site-specificity of P4ha1 in fibrillar collagens-

P4ha1 is the predominant collagen modifying enzyme, ubiquitously expressed in all tissues. A complete deletion (P4ha1−/−) of P4ha1 results in embryonically lethal mice [23,33]. Deletion of 1 allele (P4ha1+/−) retains the prolyl-4-hydroxylation activity resulting in a viable mouse. We quantitated the 4-HyP site-specific occupancy level present in different fibrillar collagen chains of wild-type and C-P4h mutants extracted from the skin. We found **23** sites on fibrillar collagen chains (Col1a1, Col1a2, and Col3a1) to be fully (≥99%) 4-hydroxylated in the wild type as well as in P4ha1 (+/−) and in different P4ha2 mutants (Table 2). Neither the occupancy levels of these specific 4-HyP sites were altered upon the complete deletion of P4ha2 nor the partial deletion of P4ha1. Additionally, these sites were also full 4-hydroxylated in wild-type where P4ha3 levels are also bare minimum. This highlights the site-specificity of P4ha1 on fibrillar collagen chains. In our analysis, these sites are considered to be P4ha1-specific sites (Table 2). The occurrence of these sites in fibrillar chains indicates the high significance of these sites in maintaining collagen structural stability and functioning in skin ECM.

**Table 2:** P4ha1 specific 4-hydroxyproline sites with occupancy (%) in mice skin fibrillar collagen chains -

| Site | Motif | WT | P4ha1$^{+/+}$;<br>P4ha2$^{+/-}$ | P4ha1$^{+/-}$;<br>P4ha2$^{+/-}$ | P4ha1$^{+/+}$;<br>P4ha2$^{-/-}$ | P4ha1$^{+/-}$;<br>P4ha2$^{-/-}$ |
| --- | --- | --- | --- | --- | --- | --- |
| Colla1 P236 | PPG | 100 | 100 | 100 | 100 | 100 |
| Colla1 P245 | LPG | 100 | 100 | 100 | 100 | 100 |
| Colla1 P305 | PPG | 100 | 100 | 100 | 100 | 100 |
| Colla1 P533 | SPG | 100 | 100 | 100 | 100 | 100 |
| Colla1 P875 | PPG | 100 | 100 | 100 | 100 | 100 |
| **Coll1a1 P629** | SPG | 100 | 100 | 100 | 100 | 100 |
| **Coll1a1 P728** | LPG | 100 | 100 | 100 | 100 | 100 |
| **Coll1a1 P860** | PPG | 99.82±0.03 | 99.71±0.15 | 99.03±0.29 | 99.89±0.19 | 99.93±0.04 |
| **Coll1a1 P866** | FPG | 99.96±0.05 | 99.93±0.07 | 99.53±0.13 | 99.89±0.20 | 99.96±0.04 |
| **Coll1a2 P174** | FPG | 100 | 100 | 100 | 100 | 100 |
| **Coll1a2 P321** | LPG | 100 | 100 | 100 | 100 | 100 |
| **Coll1a2 P327** | APG | 100 | 100 | 100 | 100 | 100 |
| **Coll1a2 P540** | PPG | 100 | 100 | 100 | 100 | 100 |
| **Coll1a2 P891** | LPG | 100 | 100 | 100 | 100 | 100 |
| **Coll1a2 P957** | YPG | 100 | 100 | 100 | 100 | 100 |
| **Coll1a2 P426** | PPG | 98.98±1.22 | 99.79±0.18 | 99.68±0.46 | 99.88±0.13 | 99.40±0.29 |
| **Col3a1 P454** | SPG | 100 | 100 | 100 | 100 | 100 |
| **Col3a1 P667** | APG | 100 | 100 | 100 | 100 | 100 |
| **Col3a1 P865** | SPG | 100 | 100 | 100 | 100 | 100 |
| **Col3a1 P868** | GPG | 100 | 100 | 100 | 100 | 100 |
| **Col3a1 P874** | FPG | 100 | 100 | 100 | 100 | 100 |
| **Col3a1 P532** | GPG | 100 | 100 | 100 | 99.85±0.09 | 99.42±0.11 |
| **Col3a1 P580** | FPG | 100 | 100 | 100 | 100 | 100 |

#### 3.4.2 Identification of site-specificity of P4ha2 in fibrillar collagens-

P4ha2 is the second abundant isoform of the C-P4h family. Although the complete deletion of P4ha2 is not lethal in mice but this collagen-modifying enzyme has a significant role in collagen deposition in the ECM. We quantitated 4-hydroxyproline occupancy levels in fibrillar collagen chains extracted from wild-type and C-P4h mutants’ mice skin. On **8** 4-HyP sites of Col1a1, Col1a2, and Col3a1, we found that there was almost full occupancy level of 4-hydroxylation on wild type, P4ha1+/+; P4ha2+/−, and P4ha1+/−; P4ha2+/−. However, 4-hydroxyproline occupancy on these sites was significantly diminished in P4ha1+/+; P4ha2−/− and P4ha1+/−; P4ha2−/− mice. This indicates that upon complete deletion of P4ha2, these sites were not modified to the levels of wild-type (Figure 4 **and** Table 3). These sites have specificity for P4ha2 enzymes. **2** P4ha2 specific sites were detected on the “-Glu-4HyP-Gly-” motif, **2** sites were detected on the -Pro-4HyP-Gly-motif, **1** site was detected on -Asn-4HyP-Gly motif, **1** site was detected on -Val-4HyP-Gly motif, and **2** sites were detected on “Thr-4HyP-Gly” motifs. These findings indicate that P4ha2 might have motif specificity in fibrillar collagens, but the motif specificity is not limited to the -Glu-4HyP-Gly motif (Table 3).

**Figure 4:**
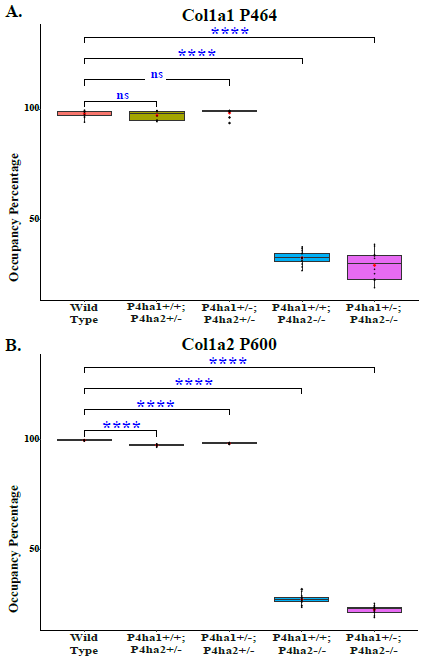
Occupancy levels of P4ha2 specific 4-HyP sites-. On the P4ha2 specific sites P464 (A) and P600 (B), the 4-hydroxyproline occupancy is almost similar in wild-type, P4ha1+/+; P4ha2+/−, and P4ha1+/−; P4ha2+/− mice but occupancy is significantly decreased in mice having complete deletion of P4ha2 (P4ha1+/+; P4ha2−/−, and P4ha1+/−; P4ha2−/−).

**Table 3:**
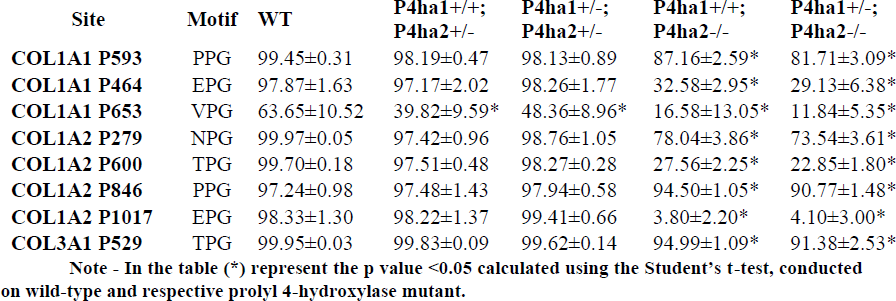
P4ha2 specific 4-hydroxyproline sites with occupancy (%) in mice skin-

| Site | Motif | WT | P4ha1+/+;<br>P4ha2+/- | P4ha1+/-;<br>P4ha2+/- | P4ha1+/+;<br>P4ha2-/- | P4ha1+/-;<br>P4ha2-/- |
| --- | --- | --- | --- | --- | --- | --- |
| **COL1A1 P593** | PPG | 99.45±0.31 | 98.19±0.47 | 98.13±0.89 | 87.16±2.59* | 81.71±3.09* |
| **COL1A1 P464** | EPG | 97.87±1.63 | 97.17±2.02 | 98.26±1.77 | 32.58±2.95* | 29.13±6.38* |
| **COL1A1 P653** | VPG | 63.65±10.52 | 39.82±9.59* | 48.36±8.96* | 16.58±13.05* | 11.84±5.35* |
| **COL1A2 P279** | NPG | 99.97±0.05 | 97.42±0.96 | 98.76±1.05 | 78.04±3.86* | 73.54±3.61* |
| **COL1A2 P600** | TPG | 99.70±0.18 | 97.51±0.48 | 98.27±0.28 | 27.56±2.25* | 22.85±1.80* |
| **COL1A2 P846** | PPG | 97.24±0.98 | 97.48±1.43 | 97.94±0.58 | 94.50±1.05* | 90.77±1.48* |
| **COL1A2 P1017** | EPG | 98.33±1.30 | 98.22±1.37 | 99.41±0.66 | 3.80±2.20* | 4.10±3.00* |
| **COL3A1 P529** | TPG | 99.95±0.03 | 99.83±0.09 | 99.62±0.14 | 94.99±1.09* | 91.38±2.53* |

Note - In the table (\*) represent the p value <0.05 calculated using the Student's t-test, conducted on wild-type and respective prolyl 4-hydroxylase mutant.

#### 3.4.3 Promiscuous sites on fibrillar collagen chains for different prolyl-4-hydroxylases

Three different isoforms of P4hs catalyze the 4-hydroxylation of proline residues present on different fibrillar collagen. Although there is site-substrate specificity for P4ha1, and P4ha2, there are certain promiscuous sites where both or all the isoforms (including P4ha3) could catalyze the 4-hydroxylation reaction. A total of 4 4-HyP sites were distinctly identified on fibrillar collagen chains, where the catalysis can occur by both P4ha1 and P4ha2 (Figure 5 and Table 4). In general, P4ha3 is the least abundant isoform of C-P4hs in wild-type tissues. However, the level of P4ha3 was found to be elevated upon partial deletion of P4ha1 and partial or complete deletion of P4ha2 in mice [52]. P4ha1+/+; P4ha2−/− and P4ha1+/−; P4ha2−/− were reported to have highly elevated mRNA levels of P4ha3 [52]. Both of these mutants have a complete deletion of P4ha2, which indicates that complete deletion of P4ha2 can highly elevate the level of P4ha3. Although the elevated level of P4ha3 upon P4ha1 and P4ha2 deletion was reported, the site-specific activity of P4ha3 upon P4ha1 and P4ha2 deletion has not been documented yet. Site-specific quantitation of the 4-hydroxyproline occupancy in mice skin fibrillar collagen chains resulted in the identification of 8 sites having decreased occupancy upon partial deletion of P4ha1 but the 4-hydroxyproline levels on these sites were not reduced upon partial deletion of P4ha2 (Table 5). The 4-hydroxyproline level on these 8 sites were either similar to wild-type or slightly elevated than the wild-type upon complete deletion of P4ha2 (Table 5). These sites were probably compensated by P4ha3 upon complete deletion of P4ha2. We detected decreased 4-hydroxyproline occupancy on 6 sites upon partial deletion of P4ha1 or P4ha2 (Table 6). However, we detected 4-hydroxyproline occupancy on these 6 sites to be with elevated in P4ha1+/+; P4ha2−/− and P4ha1+/−; P4ha2−/− mutants. We consider the elevation in the 4-hydroxyproline on these proline sites also due to the increased activity of P4ha3 upon complete P4ha2 deletion (Figure 5 and Table 6).

**Figure 5:**
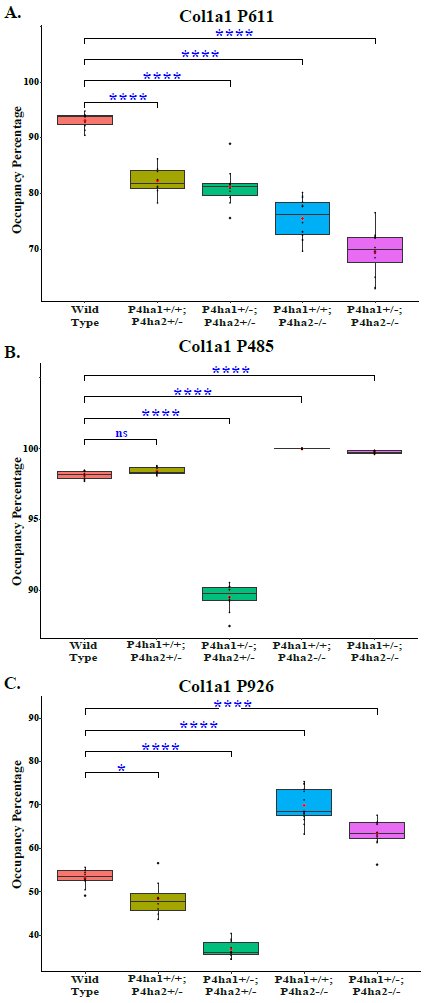
Occupancy levels of promiscuous sites for prolyl 4-hydroxylases-. (A) Col1a1 P611 site has specificity for P4ha1 and P4ha2. Partial deletion of P4ha1 and partial or complete deletion of P4ha2 significantly decrease the 4-hydroxyproline occupancy on P611 compared to the wild-type mice. (B) The 4-hydroxyproline site Col1a1 P485 has decreased occupancy P4ha1+/−; P4ha2+/− mice compared to wild-type but the 4-hydroxylation on this site is compensated by P4ha3 upon complete deletion of P4ha2 (P4ha1+/+; P4ha2−/−, and P4ha1+/−; P4ha2−/−). (C) 4-hydroxyproline occupancy on Col1a1 P926 decreased upon partial deletion of P4ha2 and P4ha1. However, 4-hydroxyproline occupancy elevates upon complete deletion of P4ha2. Increased 4-hydroxyproline occupancy on P926 can be due to elevated activity of P4ha3 upon P4ha2 deletion. This indicates that this proline site can be 4-hydroxylated with P4ha1, P4ha2, and P4ha3.

**Table 4:** P4ha1 and P4ha2 common 4-hydroxyproline sites with occupancy (%) -

| Site | Motif | WT | P4ha1<sup>+/+</sup>;<br>P4ha2<sup>+/-</sup> | P4ha1<sup>+/-</sup>;<br>P4ha2<sup>+/-</sup> | P4ha1<sup>+/+</sup>;<br>P4ha2<sup>-/-</sup> | P4ha1<sup>+/-</sup>;<br>P4ha2<sup>-/-</sup> |
| --- | --- | --- | --- | --- | --- | --- |
| COL1A1 P611 | APG | 93.11±1.31 | 82.28±2.35* | 81.21±3.32* | 75.53±3.39* | 69.52±3.93* |
| COL1A2 P789 | PPG | 94.60±0.39 | 85.43±1.54* | 84.78±2.14* | 73.83±3.44* | 65.01±3.88* |
| COL1A2 P1077 | QPG | 17.36±1.64 | 11.94±0.86* | 6.74±0.86* | 10.68±1.20* | 7.11±0.79* |
| COL3A1 P598 | GPG | 98.46±0.33 | 95.79±0.72* | 93.27±0.98* | 95.85±1.03* | 92.40±1.09* |

Note - In the table (\*) represent the p-value <0.05 calculated using the Student's t-test, conducted on wild-type and respective prolyl 4-hydroxylase mutant.

**Table 5:** P4ha1 and P4ha3 common 4-hydroxyproline sites with occupancy (%) -

| Site | Motif | WT | P4ha1<sup>+/+</sup>;<br>P4ha2<sup>+/-</sup> | P4ha1<sup>+/-</sup>;<br>P4ha2<sup>+/-</sup> | P4ha1<sup>+/+</sup>;<br>P4ha2<sup>-/-</sup> | P4ha1<sup>+/-</sup>;<br>P4ha2<sup>-/-</sup> |
| --- | --- | --- | --- | --- | --- | --- |
| Colla1 P953 | LPG | 99.47±0.09 | 99.37±0.26 | 98.04±0.61* | 99.98±0.02 | 99.96±0.03 |
| Colla1 P485 | FPG | 98.13±0.29 | 98.4±0.24 | 89.52±0.90* | 99.97±0.02 | 99.73±0.09 |
| Colla1 P680 | PPG | 97.33±0.38 | 96.28±0.53 | 89.82±1.28* | 98.75±0.29 | 97.27±0.29 |
| Colla1 P1154 | PPG | 99.73±0.06 | 99.59±0.13 | 98.11±0.23* | 99.86±0.14 | 99.74±0.03 |
| Colla2 P234 | APG | 98.99±0.48 | 95.69±3.77 | 86.39±7.70* | 99.79±0.19 | 98.37±1.28 |
| Colla2 P414 | LPG | 96.79±1.41 | 93.37±4.35 | 77.06±8.00* | 99.68±0.31 | 97.56±1.95 |
| Col3a1 P310 | RPG | 98.80±0.39 | 98.92±0.22 | 96.03±0.57* | 99.87±0.08 | 99.38±0.18 |
| Col3a1 P574 | QPG | 96.67±0.49 | 95.43±0.85 | 90.97±1.95* | 98.91±0.73 | 97.52±1.17 |

Note - In the table (\*) represent the p-value <0.05 calculated using the Student's t-test, conducted on wild-type and respective prolyl 4-hydroxylase mutant.

**Table 6:**
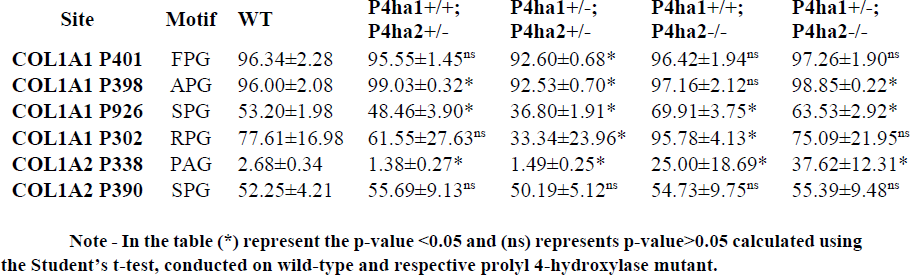
P4ha1, P4ha2 and P4ha3 common 4-hydroxyproline sites with occupancy (%) -

### 3.5 Analysis of the effect of P4ha1 and P4ha2 deletion on other collagen PTMs-

We detected P4ha1 and P4ha2 specific sites on fibrillar collagen chains from mice skin. We found that deletion of P4ha2 can diminish 4-hydroxylation on specific proline sites. But the effect of P4ha1 and P4ha2 deletion on collagen PTMs other than 4-hydroxyproline is not known till now. In order to delineate this, we analyzed the effects of partial deletion of P4ha1 and partial or complete deletion of P4ha2 on 3-hydroxyproline, 5-hydroxylysine, and O-glycosylation residues on Col1a1 and Col1a2.

#### 3.5.1.1 Analysis of the effect of P4ha1 and P4ha2 deletion on the site-specific 3-hydroxylation-

We analyzed the effect of P4ha1 and P4ha2 deletion on the occupancy levels of prolyl-3-hydroxylation sites on Col1a1 and Col1a2. P3h1 is a prominent prolyl 3-hydroxylase responsible for the site-specific prolyl 3-hydroxylation on collagen I chain [17,49,53]. In collagen 1, 3 proline sites are known to be specifically 3-hydroxylated by P3h1 activity. In Col1a1, P1153 (P986) and P874 (P707) are modified by P3h1 and in Col1a2 P803 (P707) is modified by P3h1 as shown earlier [49]. We estimated the 3-hydroxyproline occupancy on these 3 classical 3-HyP sites of collagen 1. On-site Col1a1 P1153 we detected 94.65% occupancy in wild-type mice skin in accordance with the previous findings by Pokidysheva et.al. On Col1a1 P874 and Col1a2 P803, we detected 5.7% and 13.55% occupancy respectively in wild-type mice skin. However, these Col1a1 P874 and Col1a2 P803 sites were not detected in mice skin previously by Pokidysheva et. al. in 2013 [49] in an attempt to determine P3h1-specific sites. We found that partial deletion of P4ha1 and full deletion of P4ha2 significantly increases the 3-hydroxyproline occupancy on Col1a1 P874 and Col1a2 P803 (Figure 6 **and** Table 7). On-site Col1a1 P1153, we found a significant decrease of 3-hydroxyproline occupancy in P4ha1+/−; P4ha2+/− mice. However, the 3-HyP occupancy of P1153 was significantly increased in complete deletion mutants of P4ha2 mice (Figure 6 **and** Table 7). This indicates that complete deletion of P4ha2 enhances the site-specific prolyl 3-hydroxylation on classical sites of P3h1on collagen 1.

**Figure 6:**
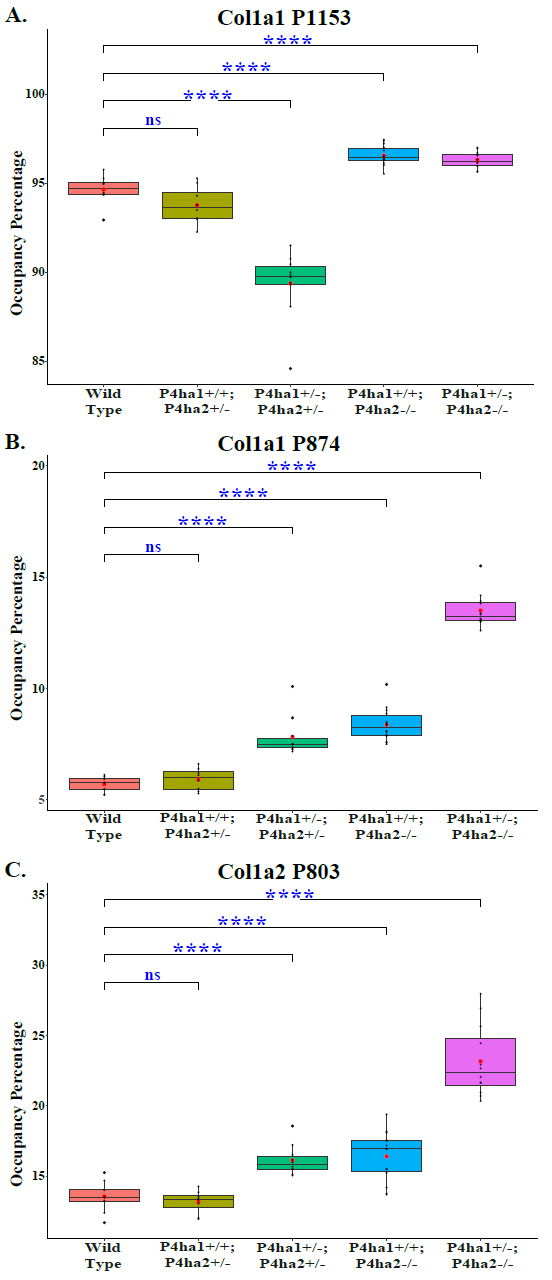
Altered occupancy levels of P3h1 specific 3-hydroxyproline sites of collagen I upon P4ha1 and P4ha2 deletion-. (A) The 3-hydroxyproline occupancy level on Col1a1 P1153 (P986) gets decreased compared to wild-type upon partial deletion of P4ha1 but it gets increased upon complete deletion of P4ha2. On the other hand, 3-hydroxyproline occupancy on (B) Col1a1 P874 (Col1a1 P707) and (C) Col1a1 P803 (P707) gets elevated compared to wild-type mice upon partial deletion of P4ha1 and/or complete deletion of P4ha2.

**Table 7:** Quantitation of 3-hydroxyproline occupancy (%) on P3h1-specific sites in collagen I.

| Site | WT | P4ha1+/+;<br>P4ha2+/- | P4ha1+/-;<br>P4ha2+/- | P4ha1+/+;<br>P4ha2-/- | P4ha1+/-;<br>P4ha2-/- |
| --- | --- | --- | --- | --- | --- |
| Colla1 P874 | 5.70±0.31 | 5.92±0.46<sup>ns</sup> | 7.83±0.86* | 8.37±0.72* | 13.51±0.74* |
| Colla1 P1153 | 94.65±0.72 | 93.77±0.98<sup>ns</sup> | 89.37±1.81* | 96.56±0.53* | 96.30±0.44* |
| Colla2 P803 | 13.55±0.98 | 13.18±0.77<sup>ns</sup> | 16.13±1.02* | 16.40±1.71* | 23.16±2.42* |

Note - In the table (\*) represent the p-value <0.05 and (ns) represents p-value>0.05 calculated using the Student's t-test, conducted on wild-type and respective prolyl 4-hydroxylase mutant.

#### 3.5.1.2 Correlation analysis of site-specific occupancy of 3-HyP1153 and 4-HyP1154 on collagen 1 alpha 1 chain-

Prolyl 3-hydroxylation on collagen chains are rare PTMs compared to prolyl-4-hydroxylation. The biological role of 3-HyP is still not well understood. The prolyl 3-hydroxylation generally occurs on the “Xaa” position of the “-Xaa-4-HyP-Gly” motif. The occurrence of 4-HyP in the “Yaa” position facilitates the occurrence of 3-HyP in the Xaa position of the repetitive triplet. This notion is substantiated by correlation analysis as represented in Figure 7. The main prolyl 3-hydroxylation site on collagen 1 alpha 1 is catalyzed by P3h1 at 3-HyP^1153^ residue. Here, we show that as the 4-hydroxylation occupancy of 4-HyP^1154^ drops in the P4ha1 +/−; P4ha2 +/− mice, the occupancy of the 3-hydroxylation site on P1153 also gets decreased. However, as the 4-hydroxylation occupancy gets complemented in the double allele deletion mutant of P4ha2, the 3-hydroxylation level increases. These results strengthen the notion that presence of 4-hydroxylation on the Yaa position facilitates the recruitment of 3-hydroxylation in the Xaa position in the repetitive sequence in collagen I chains.

**Figure 7:**
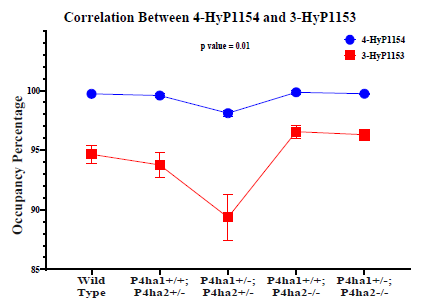
Correlation of 3-HyP1153 and 4-HyP1154 occupancy levels in Col1a1 extracted from skin of wild-type and C-P4h mutant mice-. Correlation between mice skin Col1a1 3-HyP1153 and 4-HyP1154. Correlation analysis on the occupancy levels of 3-hydroxyproline site P1153 and 4-hydroxyproline site P1154 present on -3HyP1153-4HyP1154-Gly1155-motif shows that there is a similarity in effects of partial deletion of P4ha1 and partial or complete deletion of P4ha2 on 3-HyP1153 and 4-HyP1154 site compared to the wild-type.

### 3.5.2 Analysis of the effect of P4ha1 and P4ha2 deletion on lysyl-hydroxylation -

In collagens, lysines are also heavily modified. Collagen cross-linking occurs in the ECM via the lysine residues present on the N and C terminal to form the fibrillar assembly. Hydroxylysine along with O-glycosylation are precursors for lysyl-oxidases to be oxidized to form collagen cross-link. We quantitated the hydroxylysine and O-glycosylation levels on helical collagen cross-linking sites and non-cross-linking sites. We found Col1a1 N-terminal helical cross-linking site K254 (classically known as K^87^) to be fully (∼99%) O-glycosylated (Glucosylgalactosyl-hydroxylysine) in all 5 genotypes. Thus, C-P4h mutants did not alter the classical K^87^ (Col1a1) crosslinking site. Interestingly, we found that partial deletion of P4ha1 and full deletion of P4ha2 significantly increases the hydroxylysine level in Col1a2 N-terminal helical cross-linking site K183 (Figure 8 **and** Table 8). Similarly, we found an elevation in hydroxylysine level in non-cross-linking Col1a1 K731 and Col1a2 K315 sites. We also found the occupancy level of galactosyl-hydroxylysine to be elevated upon partial deletion of P4ha1 and complete deletion of P4ha2. These findings indicate that prolyl 4-hydroxylases may crosstalk with the hydroxylation levels of helical cross-linking lysine sites and non-cross-linking lysine sites in collagen 1 (Figure 8 **and** Table 8).

**Figure 8:**
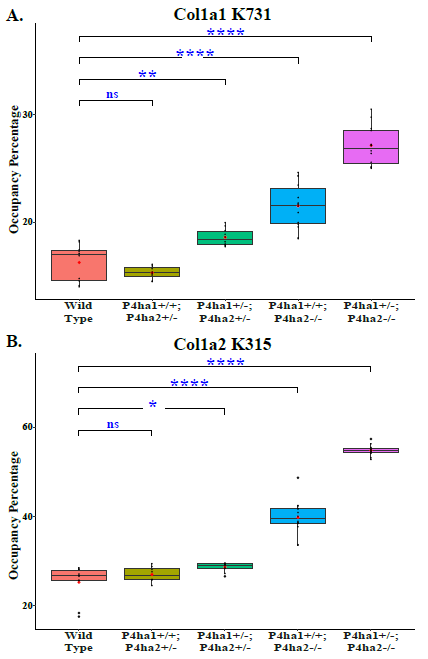
Altered occupancy levels of 2 non-cross-linking helical hydroxylysine sites upon deletion of P4ha1 and P4ha2-. Occupancy levels of collagen I helical cross-linking sites in wild-type and C-P4h mutants. 5-hydroxylysine occupancy levels on Col1a1 K731 (A) and Col1a2 K315 (B) get increased upon partial deletion of P4ha1 and/or complete deletion of P4ha2.

**Table 8:**
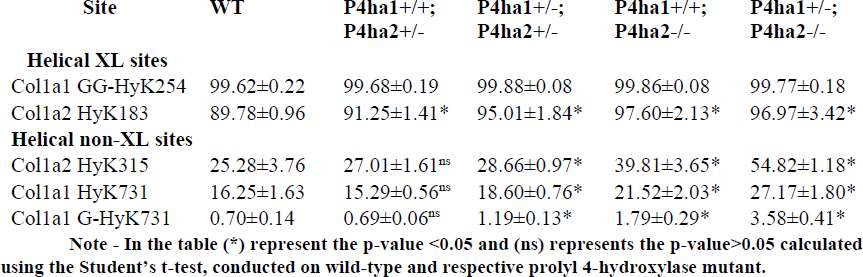
Occupancy (%) levels of lysyl hydroxylation and O-glycosylation on collagen I helical cross-linking sites and non-cross-linking sites.

## Discussion

In this study, we did a comprehensive analysis of the effects of partial deletion of P4ha1 and partial or complete deletion of P4ha2 in the deposition of collagen in the ECM. Further, the first comprehensive analysis of 4-Hyp occupancy levels, on fibrillar collagen chains (Col1a1, Col1a2, and Col3a1) was estimated. Additionally, the site-specificity of 3-HyP, 5-hydroxylysines, and O-glycosylated hydroxylysines was also determined. We found that the deletion of P4ha1 and P4ha2 in mice skin can result in the reduction in the deposition of collagen I in the skin ECM compared to the wild-type. Complete deletion of P4ha2 had a higher effect on the reduced deposition of collagen I in the ECM implicating perturbed ECM remodeling [54,55]. ECM remodeling has emerged as an important mechanism during the progression of tissue metastasis [55–57]. Elevated levels of P4ha2 are associated with poor prognosis and progression of metastasis in many cancerous tissues of the human body [56,57]. It has even been reported that inhibition/deletion of P4ha2 attenuates the metastasis progression and reduces the deposition of collagen molecules in ECM [54,55]. In our analysis, we have also found that the complete deletion of P4ha2 resulted in the reduced deposition of collagen I in the ECM.

Similarly, Tolonen at. al., [33] have also reported the significant reduction in collagen in collagen level upon complete deletion of P4ha2. Tolonen at. al., [33] performed amino acid composition analysis on collagen extracted from mice bones and we have performed label-free quantitation on MS data of collagen extracted from mice skin. But results were consistent across the tissues. Excessive deposition of collagen underlies many pathophysiological complications [5,7,55,58–63]. Therefore, P4ha2 can be a potential therapeutic target to inhibit collagen deposition in ECM in pathophysiological conditions where collagen excessively gets deposited in the ECM.

Further, we mapped the site-specific collagen PTMs in wild-type mice skin fibrillar collagen I (both alpha I and alpha II) and collagen III. Collagen I is a heterotrimer triple helix, consisting of 2 chains of Col1a1 and 1 chain of Col1a2. We identified 160 site-specific PTMs on the Col1a1 chain and we identified 124 site-specific PTMs on the Col1a2 chain. We found the Col1a2 chain to be less modified than the Col1a1 in the collagen 1 triple helix. On the other hand, collagen III is a homotrimer consisting of 3 chains of Col3a1. We mapped 137 site-specific PTMs on the Col3a1 chain. However, this chain was also found to be less modified than the Col1a1. Col1a1 chain has the highest abundance at the protein level in ECM, and the complexity of this chain is extended to harboring the highest PTMs in the wild-type mice skin fibrillar collagen chains. This analysis has yielded a detailed site-specific PTM knowledge of fibrillar collagens.

Additionally, the motif specificity of collagen prolyl 4-hydroxylases (C-P4h) was validated using high-resolution mass-spectrometry-based data. We established “-Xaa-Yaa-Gly-” as the preferred motif for prolyl-4-hydroxylation recognized by the C-P4h. We have shown the first “proteomics” based validation of the C-P4h specific motif in collagen I (Figure 3). Subsequent -Xaa-Yaa-Gly-repeats are found in the helical region of collagen 1. Repetition of this -Xaa-Yaa-Gly motifs led to the ambiguity of -Gly-Xaa-Yaa- or -Gly-Xaa-Yaa-Gly-to be the motif of C-P4h activity for prolyl-4-hydroxylation on collagen I chain. All the 4-HyP residues present on collagen I do harbor the Xaa-Yaa-Gly motif. However, because of the repetitive nature of this motif, it is difficult to delineate the exact sequence of the preferred motif. Previously, Kivirikko and Myllyharju’s group have already shown -Xaa-Yaa-Gly-as the C-P4h specific motif on collagen [2,23,24,33,44–48,64,65]. This finding is further validated by the evidence presented in this work. At the starting site of the helical region of Col1a1 and Col1a2, two specific peptides are showcased where unambiguously only Xaa-Yaa-Gly-motif is present. and 4-HyP was evidenced on the Yaa position. The -Xaa-Yaa-Gly-motif specificity for C-P4h is thus established with the high-resolution deeper proteomics analysis.

Next, the effect of partial deletion of P4ha1 and partial or complete deletion of P4ha2 on 4-hydroxyproline sites were evaluated in detail in order to delineate the site-specific substrate-specificity. We detected full 4-hydroxylation on 27 proline sites (23 novel sites reported here, Table 2) in wild type as well as C-P4h mutants. We considered these sites as P4ha1-specific sites because 4-hydroxylation on these proline sites was not diminished even upon complete deletion of P4ha2. Since P4ha3 has a lower abundance in wild-type mice tissues [33,66], it is likely that even one allele of P4ha1 (+/−) mutants was able to fully restore the complete occupancy of these sites. So, these sites can even be modified by a single allele of P4ha1 (P4ha1+/−). These sites are detected in 3 chains (Col1a1, Col1a2, and Col3a1) of fibrillar collagen (Table 2). We also found Col1a1P230 on GRPGER, Col1a1 P671 on GFPGER, Col1a1P719 on GMPGER, and Col1a2 P159 on GRPGER to be fully modified as described by Sipilä et. al., 2018 [23]. Full occupancy of 4-hydroxylation on these proline sites indicates that these sites have a high significance in collagen triple helix formation, triple helix stability, and function of fibrillar collagens. Our analysis also identified P4ha2-specific sites in collagen I. On 8 proline sites in collagen I, we found 4-hydroxylation occupancy to be diminished upon complete deletion of P4ha2. However, 4-hydroxylation occupancy levels on these sites were ∼100% in partial P4ha1 and partial P4ha2 deletion mutants. This means that these sites have specificity for P4ha2, and these sites cannot be fully modified in the complete absence of the P4ha2 enzyme. These P4ha2 specific sites have ∼100% 4-hydroxylation occupancy in wild type, so these sites might have some role in collagen biosynthesis and collagen interactions with extracellular proteins and cell-surface receptors. Recently Wilhelm et.al [67] pointed out GDP/GEP motifs as a substrate for P4ha2-specificity. However, our analysis not only identified novel sites of P4ha2 specificity but also establishes that the catalysis is not specific to GDP/GEP motifs (Table 3). Elucidation of the site-specific role of these sites in collagen biosynthesis and collagen interactions is an opportunity to explore in the future.

Moreover, in this study, we have shown for the first time that the deletion of collagen prolyl 4-hydroxylases increases the site-specific 3-hydroxyproline occupancy. We detected elevated levels of 3-hydroxyproline occupancy in Col1a1 P874, Col1a1 P1153, and Col1a2 P803 sites in mice skin upon complete deletion of P4ha2 (Figure 6). These 3-hydroxyprolines are reported to have specificity for P3h1 enzyme in mice bone collagen 1 [49]. Elevation of 3-hydroxylation on these sites indicates a possible elevated activity of P3h1 upon partial deletion of P4ha1 and complete deletion of P4ha2. These findings show that there is a strong association of P4ha2 with prolyl 3-hydroxylation activity. However, we found that there is complexity in the connection of prolyl 4-hydroxylases deletion and 3-hydroxylase activity on these 3 proline sites. The 3-hydroxylation on Col1a1 P874 and Col1a2 P803 was not changed upon partial deletion of P4ha2 (P4ha1+/+; P4ha2+/−) compared to wild-type. We identify these two sites of 3-HyP from mice skin collagen for the first time. In a previous MS analysis of collagens from the mice skin collagen did not identify these two sites. This is probably because of the lower occupancy of 3-hydroxyproline on these sites and also the limited capabilities of the mass-spectrometry-based PTM analysis pipeline. These sites are popularly known as A3 sites in Col1a1 and Col1a2. The 3-hydroxylation on Col1a1 P874 and Col1a2 P803 was elevated on the partial deletion of P4ha1 and P4ha2 (P4ha1+/−; P4ha2+/−) and 3-hydroxylation was also increased in mice having a complete deletion of P4ha2 (P4ha1+/+; P4ha2−/−, and P4ha1+/−; P4ha2−/−) compared to the wild-type mice. Among all genotypes, mice having partial deletion P4ha1 and complete deletion of P4ha2 (P4ha1+/−; P4ha2−/−) have the highest levels of 3-hydroxyproline occupancy on Col1a1 P874 and Col1a2 P803 sites. This means that partial deletion of P4ha2 cannot alter the 3-hydroxylation activity on Col1a1 P874 and Col1a2 P803 sites in mice skin but partial deletion P4ha1 or complete deletion P4ha2 can elevate the 3-hydroxylation occupancy on these 2 sites. On the other hand, there are different effects of P4ha1 deletion and P4ha2 deletion on the 3-hydroxyproline occupancy level on the Col1a1 P1153 site. This P1153 site popularly known as the A1 site, is associated with the osteogenesis imperfecta. This site has been reported to have specificity for P3h1 in mice tendon, bone, and skin tissues [49]. In our analysis, we found that the 3-hydroxylation occupancy level on this site has a slight (statistically non-significant) decrease upon partial deletion of P4ha2 (P4ha1+/+; P4ha2+/−) (Figure 6 and Table 7). But 3-hydroxylation on this site was elevated (p<0.01) in P4ha1+/+; P4ha2−/− mice. Partial deletion of P4ha1 has a different effect on P1153 than it has on Col1a1 P874 and Col1a2 P803 sites. Partial deletion of P4ha1 with partial deletion P4ha2 (P4ha1+/−; P4ha2+/−) results in a significant decrease in the 3-hydroxyproline occupancy level on the P1153 site. Even P4ha1+/−; P4ha2−/− mice have lower has lower 3-hydroxylation level than the P4ha1+/+; P4ha2−/− mice (Figure 6). This indicates that partial deletion of P4ha1 can decrease the 3-hydroxylation level on Col1a1 P1153. This analysis supported the notion of the occurrence of 3-HyP in the Xaa position of the Xaa-4-HyP-Gly motif. The level of 4-HyP on the Yaa position determines the 3-hydroxylation catalysis in the Yaa position. However, complete deletion of P4ha2 can elevate the 3-hydroxylation on P1153 compared to the wild-type. The different effects of P4ha1 deletion and P4ha2 deletion on this hints towards different modes of interaction of P4ha1 and P4ha2 on prolyl 3-hydroxylases.

Our analysis, for the first-time highlights that the effect of prolyl 4-hydroxylase deletion is not limited to only the proline modifications. Here, we show that lysine modifications are also affected by P4ha1 and P4ha2 deletion. Despite having enzymatic activities of 4-hydroxylation on proline, P4ha1 and P4ha2 deletion can affect the lysine modifications in a site-specific manner. We found significantly elevated levels of 5-hydroxylysine occupancy on the collagen 1 alpha 2 helical cross-linking lysine site (Col1A2 K^183^). Modulation in 5-hydroxylysine occupancy on helical cross-linking site upon P4ha2 deletion may have an effect on collagen cross-linking and subsequent fibrillar assembly. However, the O-glycosylation (glucosyl-galactosyl-hydroxylysine form) level of classical helical crosslinking site α1K^87^ (referred to here as Col1A1 K^254^) was not affected upon deletion of C-P4hs. Interestingly, >99% of this K^254^ site is in glucosylgalactosyl-hydroxylysine form with almost no microheterogeneity. The same site in bovine skin has been shown to harbor microheterogeneity with galactosyl-hydroxylysine form as the most abundant one, however, the same site in mice bone was present in abundance mostly with the disaccharide form of O-glycosylation [68]. The evaluation of non-cross-linking lysine sites in mice having a partial deletion of P4ha1 (P4ha1+/−; P4ha2+/− and (P4ha1+/−; P4ha2−/−) and mice having a complete deletion of P4ha2 (P4ha1+/+; P4ha2−/− and P4ha1+/−; P4ha2−/−) compared to the wild-type mice revealed significantly increased hydroxylation and galactosyl-hydroxylation on Col1a1 K^731^ site (Figure 8 & Table 8). We found that partial deletion of P4ha1 and complete deletion of P4ha2 can elevate the 5-hydroxylysine occupancy level on helical cross-linking and non-cross-linking lysine sites. These findings indicate that P4ha1 and P4ha2 can modulate the activity of lysyl-hydroxylases on collagen. Collagen cross-linking analysis on P4ha2 deleted or overexpressed mice would delineate the exact effects of P4ha2 on collagen cross-linking and the mechanism of fibrillar assembly. This study hints towards the crosstalk between prolyl 4-hydroxylases and lysyl hydroxylases. It will be interesting to explore the possibility of direct interaction of prolyl 4-hydroxylases with lysyl hydroxylases and also the role of their interaction in collagen biosynthesis and the functioning of collagen chains.

## Author contributions

**VS**: Conceptualization, Methodology, Data curation, Software, Formal analysis, Investigation, Visualization, Writing original draft; **TB:** Conceptualization, Supervision, Review & editing.

## Acknowledgments and data availability

The authors acknowledge Prof. Jyrki Heino and Dr. Pekka Rappu for publicly sharing the raw mass spectrometry dataset (PXD008802) [23] on ProteomeXchange Consortium via the PRIDE partner repository. All other analyzed files can be shared upon a valid request to VS.

