## Supplementary material for "Decoding the comprehensive substrate-specificity and evidence of altered site-specific collagen prolyl-3-hydroxylation, lysyl-hydroxylation, and lysyl O-glycosylation in P4ha1 and P4ha2 deleted mutant mice": Sarohi MB Supporting Information.pdf

**Table of Contents**


| S.N. | Title | Page No. |
|---|---|---|
| 1 | Supplementary Table 1 | 2 |
| 2 | Supplementary Figure 1(A) (4-HyP sites) | 3-6 |
| 3 | Supplementary Figure 1(B) (3-HyP sites) | 7-9 |
| 4 | Supplementary Figure 1(C) (HyK sites) | 10-11 |

**Supplementary Table 1:** Relative expression of fibrillar collagen chains in wild-type and prolyl 4-hydroxylase mutant mice. In the table (*) represent the p value <0.05 from the Student's t-test conducted on wild-type and respective prolyl 4-hydroxylase mutant.

| Collagen Chains | WT | P4ha1+/+; P4ha2+/- | P4ha1+/-; P4ha2+/- | P4ha1+/+; P4ha2-/- | P4ha1+/-; P4ha2-/- |
|---|---|---|---|---|---|
| **Col1a1** | 35.02±0.08 | 34.86±0.06* | 34.85±0.07* | 34.78±0.24* | 34.76±0.11* |
| **Col1a2** | 33.93±0.10 | 33.79±0.05* | 33.75±0.05* | 33.62±0.23* | 33.55±0.15* |
| **Col3a1** | 33.38±0.19 | $33.46±0.26^{ns}$ | $33.34±0.13^{ns}$ | $33.30±0.24^{ns}$ | 33.81±0.16* |

**Supplementary Figure 1:** We identified site-specific collagen post-translational modifications in fibrillar collagen chains (Col1a1, Col1a2, and Col3a1) extracted from wild-type mice skin. Manual analysis of peptide spectrum match (PSM) was performed. PSMs with mass error lower than ±20 ppm and less than 1% FDR were considered for manual analysis. PSMs were annotated using pLabel. Supplementary figure 1(A) (page 3-6) shows representative PSMs for 4-hydroxyproline sites. Supplementary figure 1(B) (page 7-9) shows representative PSMs for 3-hydroxyproline sites and Supplementary figure 1(C) (page 10-11) shows representative PSMs for hydroxylysine sites.

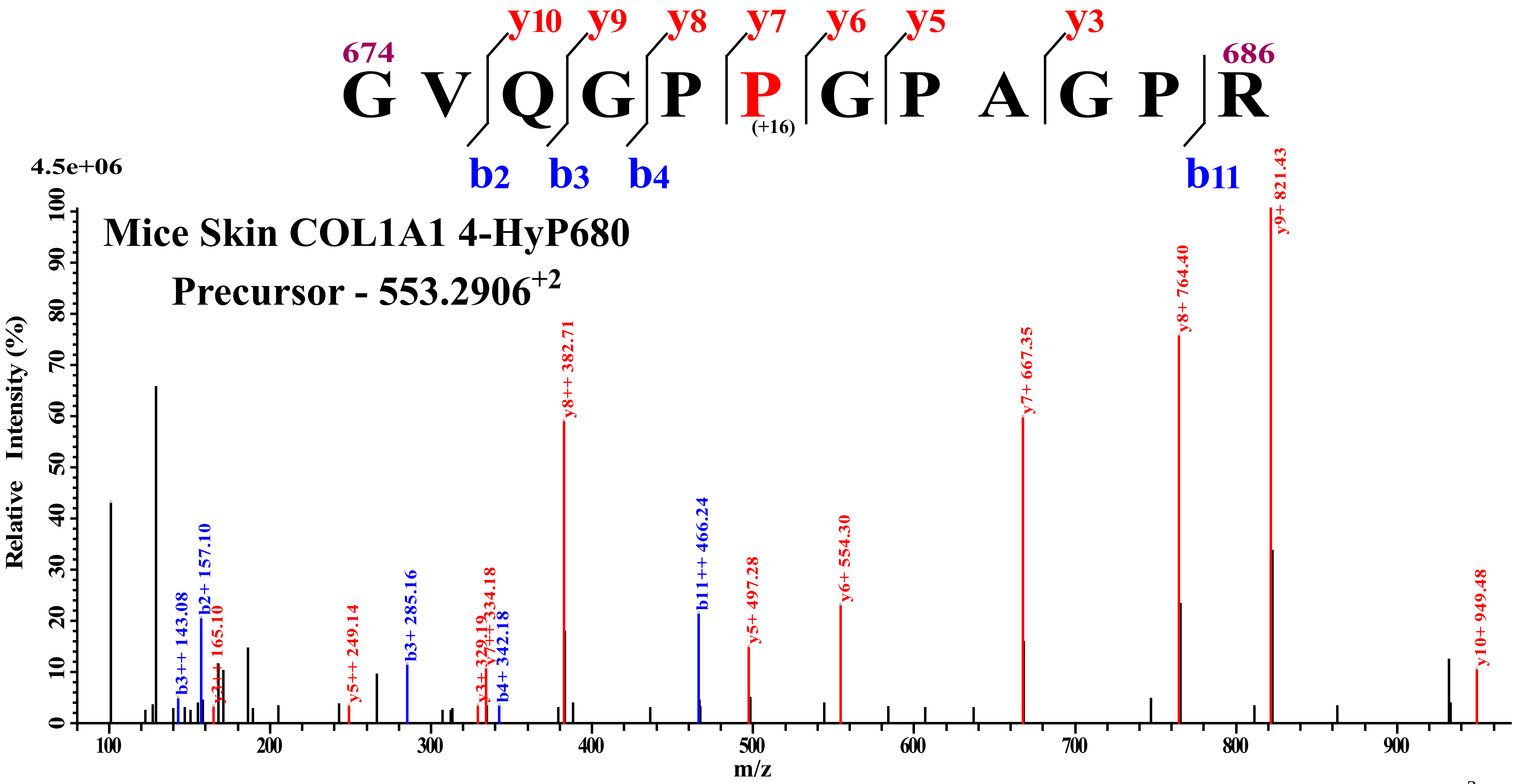

674
686
G V Q G P P G P A G P R
(+16)
y10 y9 y8 y7 y6 y5 y3
b2 b3 b4 b11
4.5e+06
Mice Skin COL1A1 4-HyP680
Precursor - 553.2906+2
Relative Intensity (%)
m/z
b3++ 143.08
b2+ 157.10
y3++ 165.10
y5++ 249.14
b3+ 285.16
y3+ 329.19
y7++ 334.18
b4+ 342.18
y8++ 382.71
b11++ 466.24
y5+ 497.28
y6+ 554.30
y7+ 667.35
y8+ 764.40
y9+ 821.43
y10+ 949.48

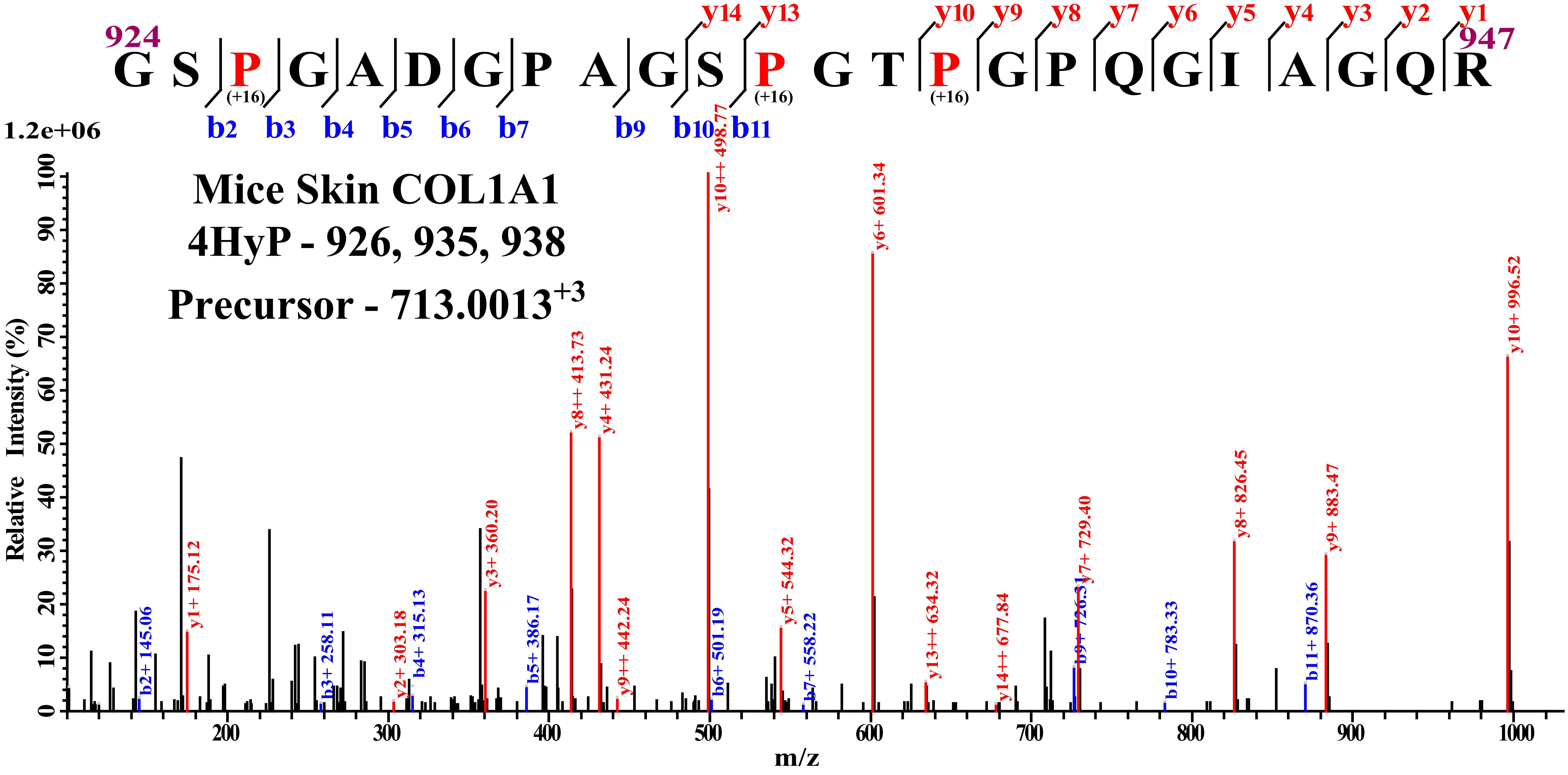

924
G S P G A D G P A G S P G T P G P Q G I A G Q R
947
(+16)
y14 y13 y10 y9 y8 y7 y6 y5 y4 y3 y2 y1
b2 b3 b4 b5 b6 b7 b9 b10 b11
1.2e+06
Mice Skin COL1A1
4HyP - 926, 935, 938
Precursor - 713.0013+3
Relative Intensity (%)
m/z
y10++ 498.77
y6+ 601.34
y10+ 996.52
y8++ 413.73
y4+ 431.24
y8+ 826.45
y9+ 883.47
y7+ 729.40
y3+ 360.20
y5+ 544.32
y13++ 634.32
y14++ 677.84
y9++ 442.24
y2+ 303.18
y1+ 175.12
b2+ 145.06
b3+ 258.11
b4+ 315.13
b5+ 386.17
b6+ 501.19
b7+ 558.22
b10+ 783.33
b11+ 870.36

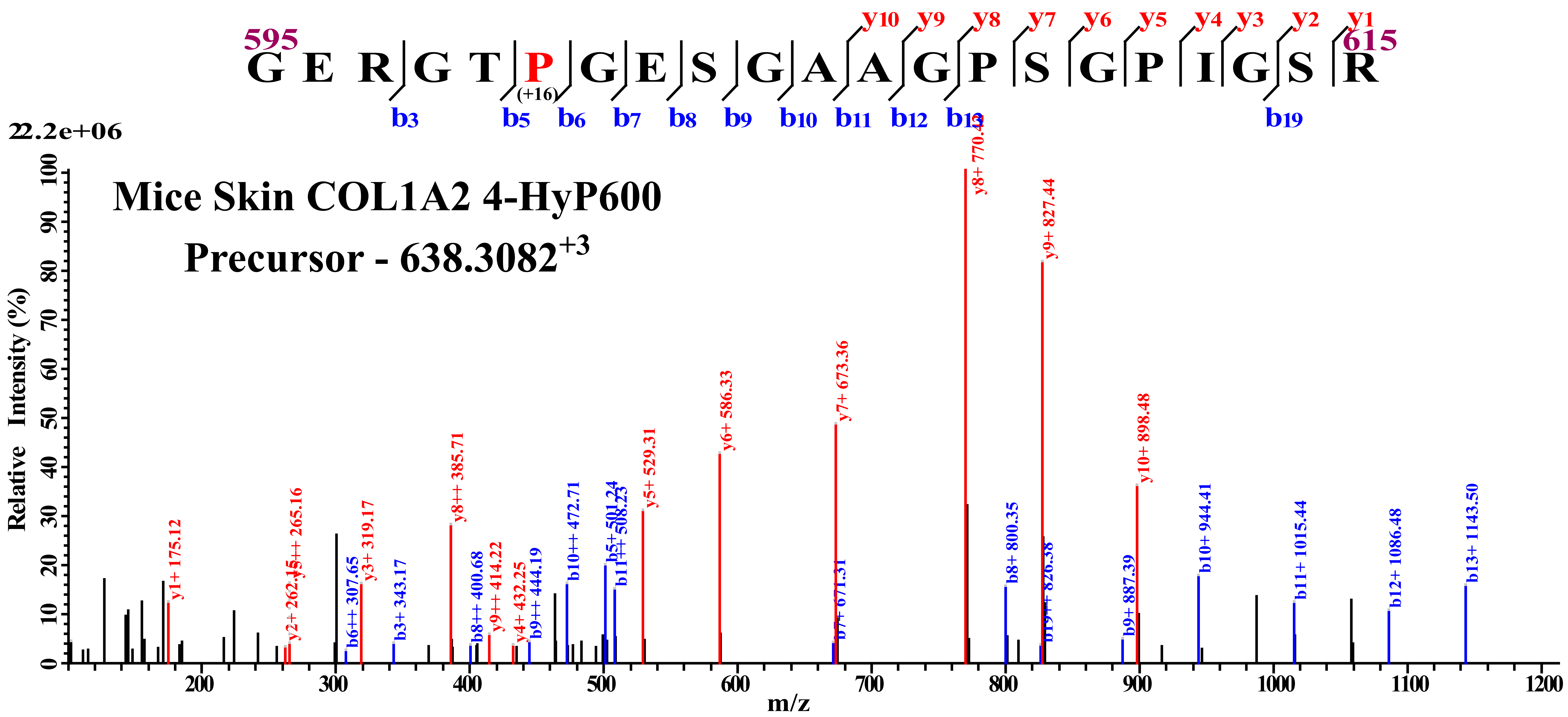

Mice Skin COL1A2 4-HyP600
Precursor - 638.3082+3
595
G E R G T P G E S G A A G P S G P I G S R
615
(+16)
b3 b5 b6 b7 b8 b9 b10 b11 b12 b13 b19
y10 y9 y8 y7 y6 y5 y4 y3 y2 y1
2.2e+06
Relative Intensity (%)
m/z
y8+ 770.43
y9+ 827.44
y7+ 673.36
y6+ 586.33
y10+ 898.48
y8++ 385.71
y5+ 529.31
b10+ 944.41
b13+ 1143.50
b10++ 472.71
b5+ 501.24
b11++ 508.23
b8+ 800.35
b11+ 1015.44
b12+ 1086.48
y1+ 175.12
y3++ 265.16
y2+ 262.15
y3+ 319.17
b6++ 307.65
b3+ 343.17
b8++ 400.68
y9++ 414.22
y4+ 432.25
b9++ 444.19
b7+ 671.31
b19++ 826.38
b9+ 887.39

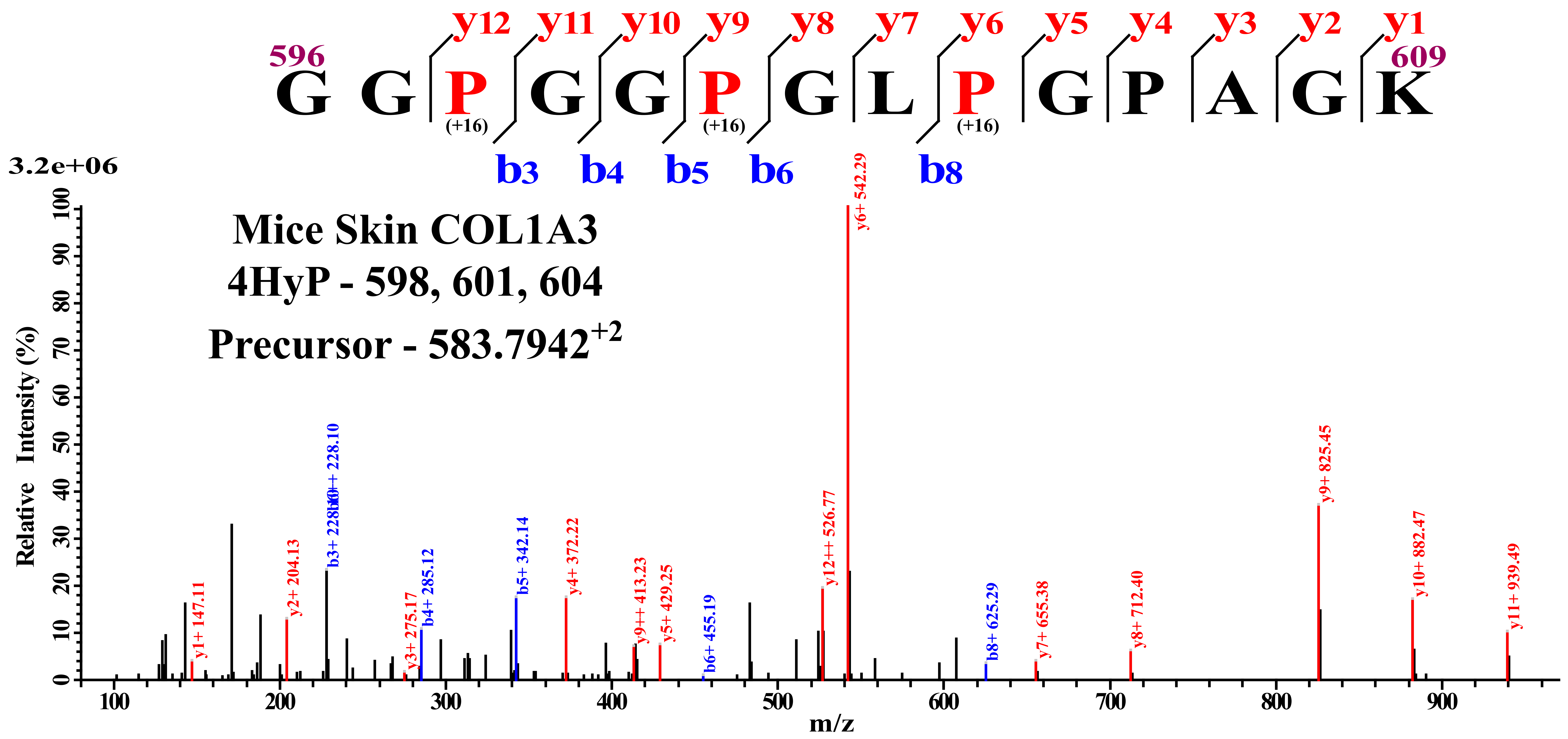

y12 y11 y10 y9 y8 y7 y6 y5 y4 y3 y2 y1
596
G G P G G P G L P G P A G K
609
(+16) (+16) (+16)
b3 b4 b5 b6 b8
3.2e+06
Mice Skin COL1A3
4HyP - 598, 601, 604
Precursor - 583.7942+2
Relative Intensity (%)
m/z
y6+ 542.29
y9+ 825.45
y12++ 526.77
b3+ 228.10
y10+ 882.47
y11+ 939.49
y1+ 147.11
y2+ 204.13
y3+ 275.17
b4+ 285.12
b5+ 342.14
y4+ 372.22
y9++ 413.23
y5+ 429.25
b6+ 455.19
b8+ 625.29
y7+ 655.38
y8+ 712.40

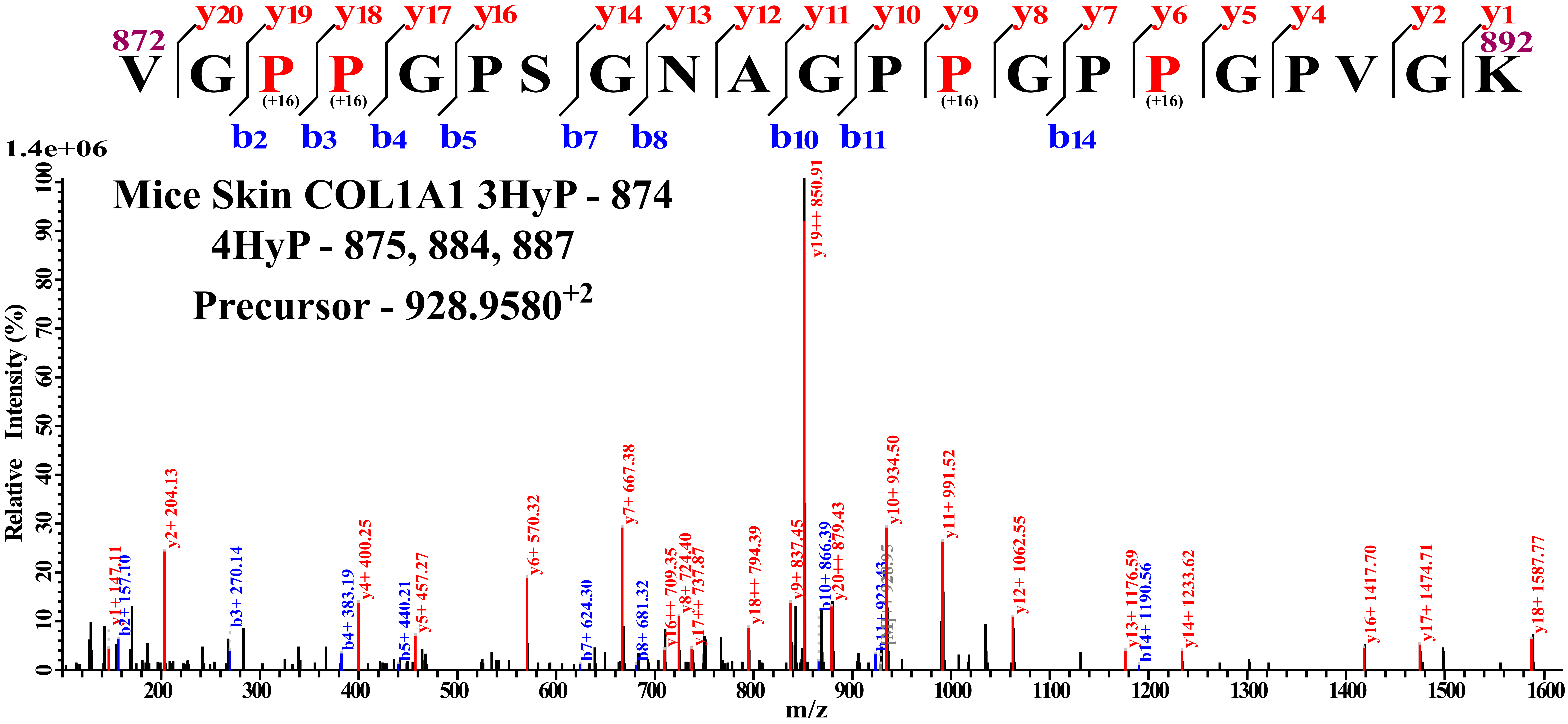

Mice Skin COL1A1 3HyP - 874
4HyP - 875, 884, 887
Precursor - 928.9580+2
872
V G P P G P S G N A G P P G P P G P V G K
892
(+16) (+16) (+16) (+16)
y20 y19 y18 y17 y16 y14 y13 y12 y11 y10 y9 y8 y7 y6 y5 y4 y2 y1
b2 b3 b4 b5 b7 b8 b10 b11 b14
1.4e+06
Relative Intensity (%)
m/z
y1+ 147.11
b2+ 157.10
y2+ 204.13
b3+ 270.14
b4+ 383.19
y4+ 400.25
b5+ 440.21
y5+ 457.27
y6+ 570.32
b7+ 624.30
y7+ 667.38
b8+ 681.32
y16++ 709.35
y8+ 724.40
y17++ 737.87
y18++ 794.39
y9+ 837.45
y19++ 850.91
b10+ 866.39
y20++ 879.43
b11+ 923.43
y10+ 934.50
y11+ 991.52
y12+ 1062.55
y13+ 1176.59
b14+ 1190.56
y14+ 1233.62
y16+ 1417.70
y17+ 1474.71
y18+ 1587.77

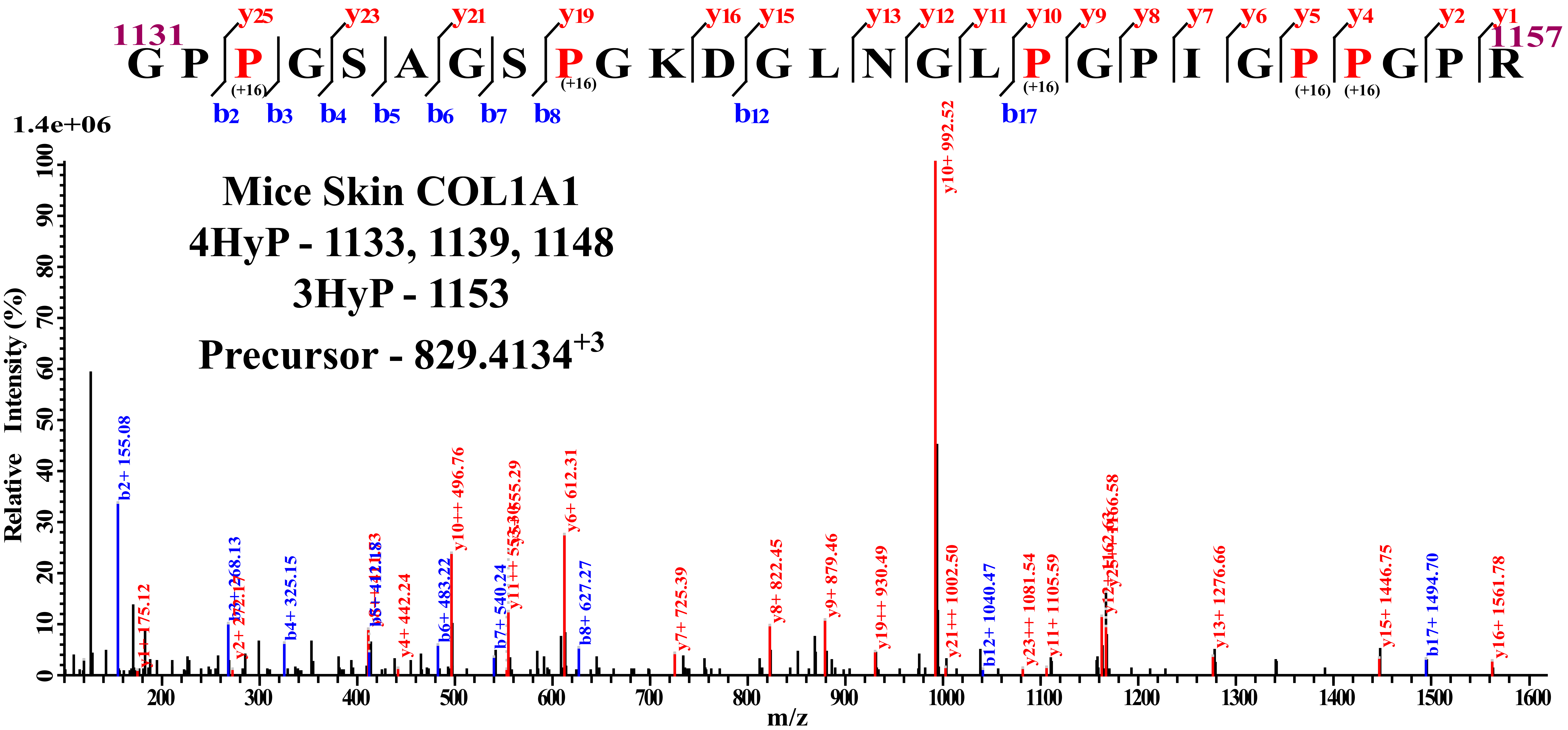
Mice Skin COL1A1
4HyP - 1133, 1139, 1148
3HyP - 1153
Precursor - 829.4134+3
1131 G P P G S A G S P G K D G L N G L P G P I G P P G P R 1157
(+16) (+16) (+16) (+16) (+16)
y25 y23 y21 y19 y16 y15 y13 y12 y11 y10 y9 y8 y7 y6 y5 y4 y2 y1
b2 b3 b4 b5 b6 b7 b8 b12 b17
1.4e+06
Relative Intensity (%)
m/z
b2+ 155.08
y1+ 175.12
b3+ 268.13
y2+ 272.17
b4+ 325.15
y4+ 442.24
b6+ 483.22
y10++ 496.76
b7+ 540.24
y11++ 555.29
y6+ 612.31
b8+ 627.27
y7+ 725.39
y8+ 822.45
y9+ 879.46
y19++ 930.49
y10+ 992.52
y21++ 1002.50
b12+ 1040.47
y23++ 1081.54
y11+ 1105.59
y12+ 1166.58
y13+ 1276.66
y15+ 1446.75
b17+ 1494.70
y16+ 1561.78

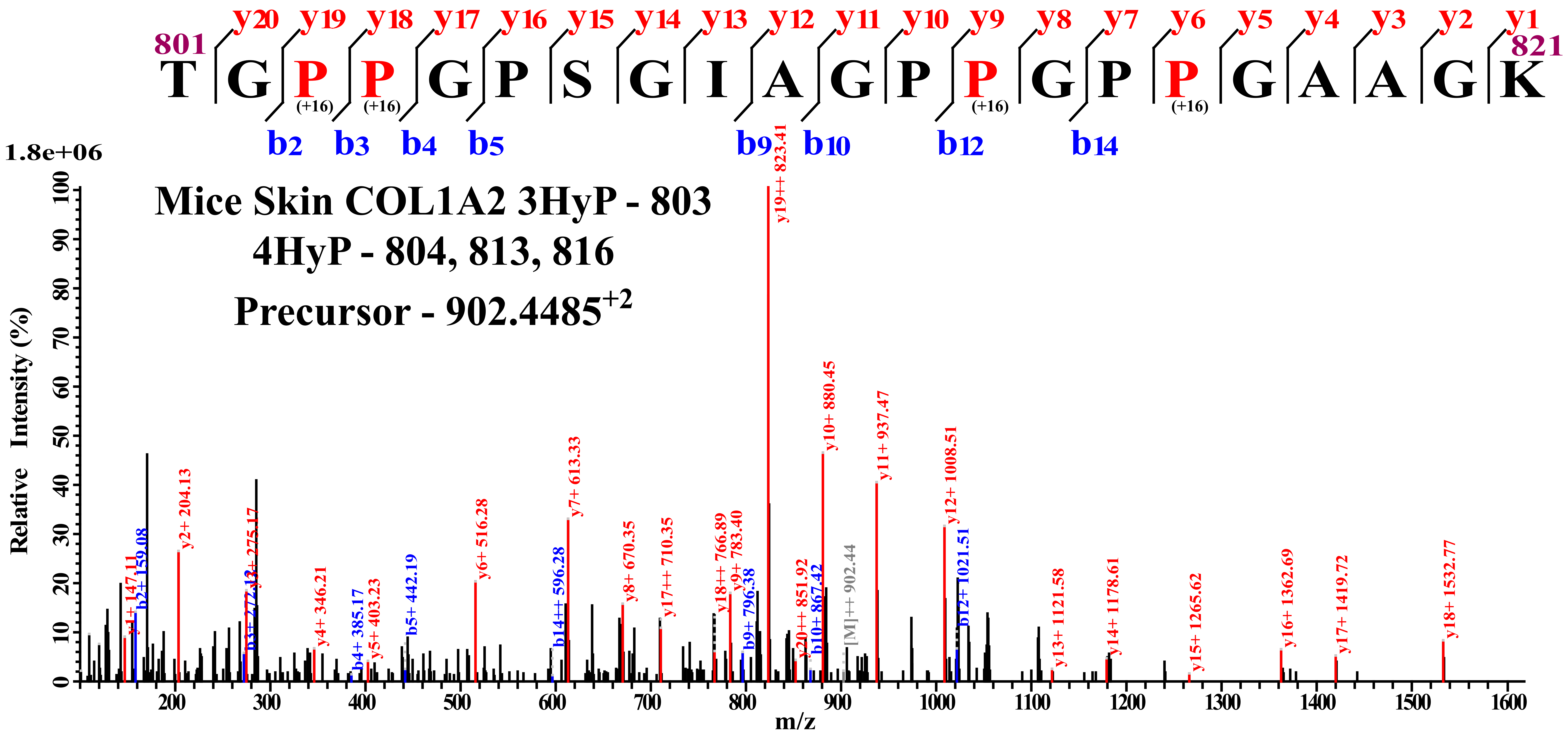

801
T G P P G P S G I A G P P G P P G A A G K
821
(+16) (+16) (+16) (+16)
y20 y19 y18 y17 y16 y15 y14 y13 y12 y11 y10 y9 y8 y7 y6 y5 y4 y3 y2 y1
b2 b3 b4 b5 b9 b10 b12 b14
Mice Skin COL1A2 3HyP - 803
4HyP - 804, 813, 816
Precursor - 902.4485+2
1.8e+06
Relative Intensity (%)
m/z
y19++ 823.41
y10+ 880.45
y11+ 937.47
y12+ 1008.51
y7+ 613.33
y2+ 204.13
b12+ 1021.51
y6+ 516.28
y1+ 147.11
b2+ 159.08
y3+ 275.17
b3+ 272.13
y4+ 346.21
b4+ 385.17
y5+ 403.23
b5+ 442.19
b14++ 596.28
y8+ 670.35
y17++ 710.35
y18++ 766.89
y9+ 783.40
b9+ 796.38
y20++ 851.92
b10+ 867.42
[M]++ 902.44
y13+ 1121.58
y14+ 1178.61
y15+ 1265.62
y16+ 1362.69
y17+ 1419.72
y18+ 1532.77

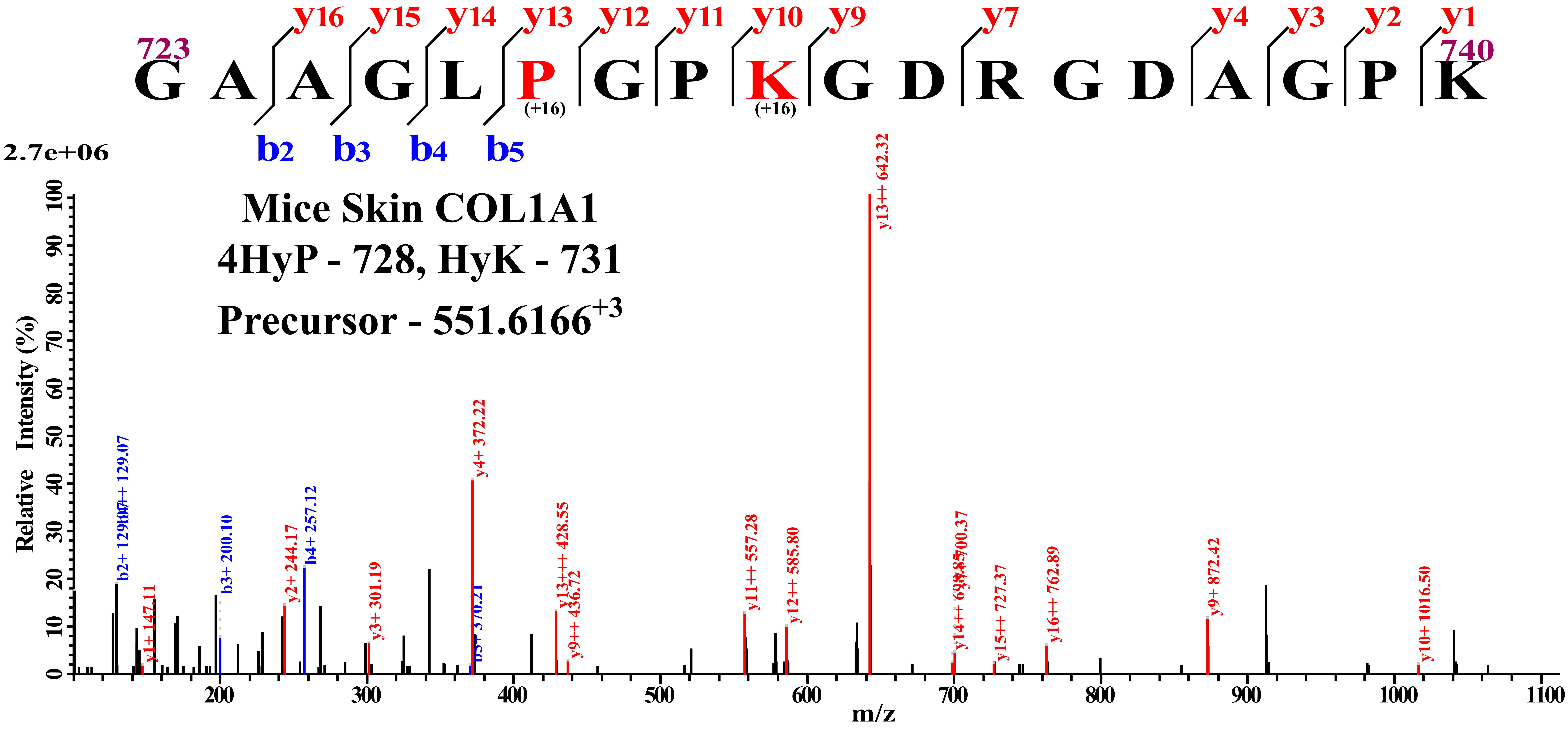
y16 y15 y14 y13 y12 y11 y10 y9 y7 y4 y3 y2 y1
723
740
G A A G L P G P K G D R G D A G P K
(+16)
(+16)
b2 b3 b4 b5
2.7e+06
Mice Skin COL1A1
4HyP - 728, HyK - 731
Precursor - 551.6166+3
Relative Intensity (%)
m/z
b2+ 129.07
b3+ 200.10
b4+ 257.12
y1+ 147.11
y2+ 244.17
y3+ 301.19
y4+ 372.22
b5+ 370.21
y13+++ 428.55
y9++ 436.72
y11++ 557.28
y12++ 585.80
y13++ 642.32
y15++ 727.37
y16++ 762.89
y9+ 872.42
y10+ 1016.50

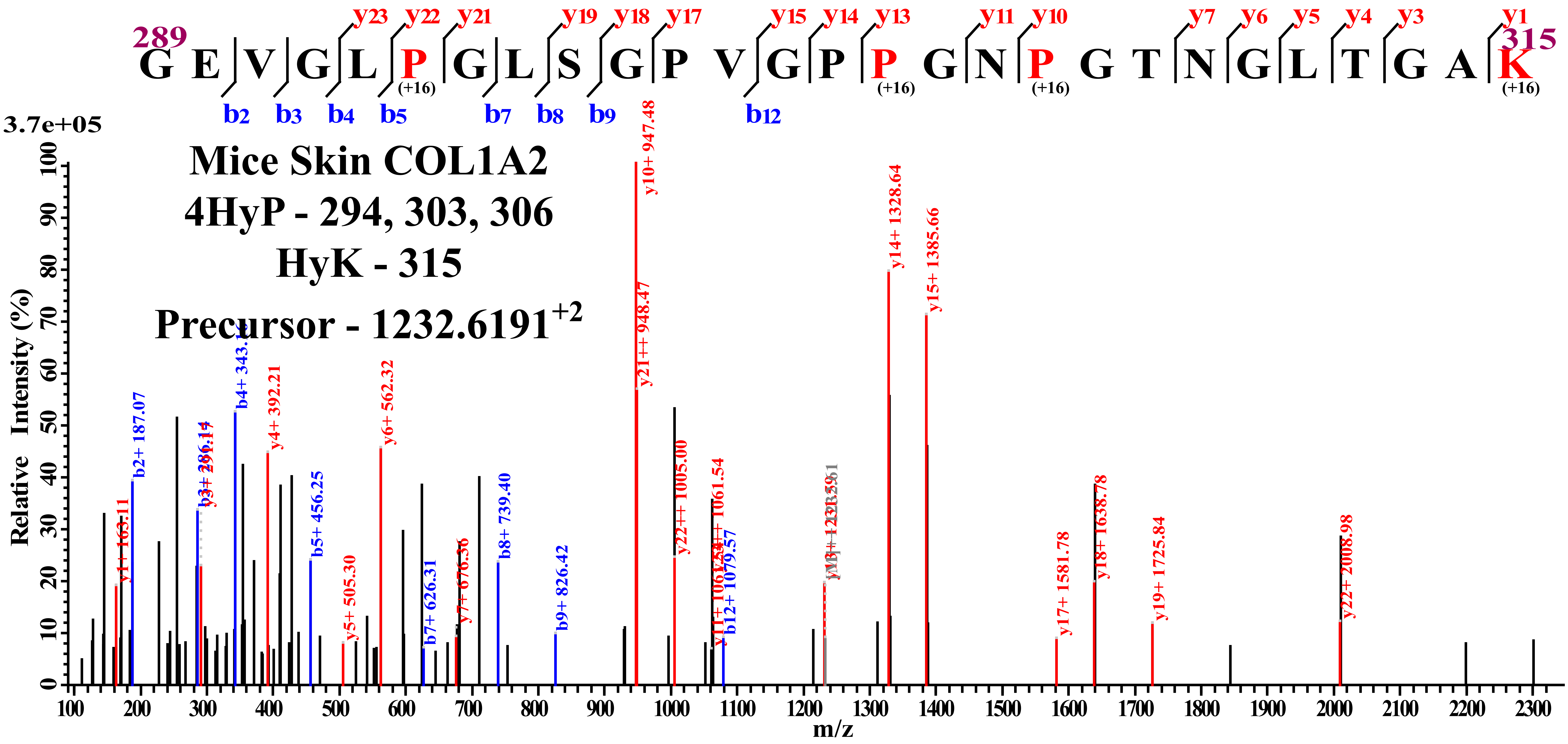
Mice Skin COL1A2
4HyP - 294, 303, 306
HyK - 315
Precursor - 1232.6191+2
289
G E V G L P G L S G P V G P P G N P G T N G L T G A K
315
(+16)
(+16)
(+16)
(+16)
y23 y22 y21 y19 y18 y17 y15 y14 y13 y11 y10 y7 y6 y5 y4 y3 y1
b2 b3 b4 b5 b7 b8 b9 b12
3.7e+05
Relative Intensity (%)
m/z
y1+ 163.11
b2+ 187.07
b3+ 286.14
y3+ 291.17
b4+ 343.16
y4+ 392.21
b5+ 456.25
y5+ 505.30
y6+ 562.32
b7+ 626.31
y7+ 676.36
b8+ 739.40
b9+ 826.42
y10+ 947.48
y21++ 948.47
y22++ 1005.00
y11+ 1061.54
b12+ 1079.57
y14+ 1328.64
y15+ 1385.66
y17+ 1581.78
y18+ 1638.78
y19+ 1725.84
y22+ 2008.98
